# Neural Anatomy and Optical Microscopy (NAOMi) Simulation for evaluating calcium imaging methods

**DOI:** 10.1101/726174

**Authors:** Adam S. Charles, Alexander Song, Jeff L. Gauthier, Jonathan W. Pillow, David W. Tank

## Abstract

The past decade has seen a multitude of new *in vivo* functional imaging methodologies. However, the lack of ground-truth comparisons or evaluation metrics makes large-scale, systematic validation impossible. Here we provide a new framework for evaluating TPM methods via *in silico* Neural Anatomy and Optical Microscopy (NAOMi) simulation. Our computationally efficient model generates large anatomical volumes of mouse cortex, simulates neural activity, and incorporates optical propagation and scanning to create realistic calcium imaging datasets. We verify NAOMi simulations against *in vivo* two-photon recordings from mouse cortex. We leverage this access to *in silico* ground truth to perform direct comparisons between different segmentation algorithms and optical designs. We find modern segmentation algorithms extract strong neural time-courses comparable to estimation using oracle spatial information, but with an increase in the false positive rate. Comparison between optical setups demonstrate improved resilience to motion artifacts in sparsely labeled samples using Bessel beams, increased signal-to-noise ratio and cell-count using low numerical aperture Gaussian beams and nuclear GCaMP, and more uniform spatial sampling with temporal focusing versus multi-plane imaging. Overall, by leveraging the rich accumulated knowledge of neural anatomy and optical physics, we provide a powerful new tool to assess and develop important methods in neural imaging.

## 1 Introduction

The endeavor to understand neural systems has spurred rapid development of technology that can record brain activity at ever larger scales [1–3] and higher precision [4–6]. One such class of technology, functional optical microscopy, has empowered researchers to explore neural dynamics from synapse [7, 8] to large brain regions [9, 10]. Specifically, two-photon microscopy (TPM) combined with in vivo calcium imaging [11–16], has enabled the simultaneous recording of unprecedented numbers of neurons (over 9000) at cellular resolution [17, 18].

Although TPM has found widespread use [19–21], many available experimental techniques and data processing algorithms lack appropriate, systematic assessment [22, 23]. This deficit can result in inaccurate interpretation of neural data [24]. A systematic comparison of techniques would allow researchers to make better informed decisions about equipment and data-processing.

For instance, while imaging deeper into scattering tissue with TPM can benefit from decreasing the excitation numerical aperture (NA) [15], it is unknown how this benefit interacts with other optical or experimental design choices, such as adaptive optics [25, 26] or dendritic imaging [27, 28]. Additionally, while many algorithms have been designed to extract the neural activity traces and spatial profiles from TPM data [29–43], few options exist to assess the fidelity of the inferred segmentation beyond comparisons to manually annotated data [24, 44, 45].

In both cases assessment suffers from a lack of ground truth data, the gold standard of which requires simultaneous intracellular electrophysiological and TPM recordings [46, 47]. Such experiments are both difficult to perform and limited to only a few neurons and imaging conditions. The small number of neurons from such experiments limits the assessment scope by biasing towards cells that are in focus, fire often, and fluoresce brightly. This problem is further exacerbated as assessing multiple imaging parameters requires recordings under each imaging condition, greatly increasing the cost of collecting such data.

Alternatively, subjective ground truth can be obtained from TPM recordings via manual annotation [48]. Human labels, however, do not provide access to the underlying neural spiking, are limited by the same signal-to-noise ratio (SNR) that limit demixing algorithms, and may also bias analysis against dim or sparsely firing neurons. These same issues also affect comparisons using simultaneous conventional TPM recordings to test novel imaging conditions [49, 50].

In place of collecting ground truth data, simulations can provide rich, controlled testing data. Such approaches have benefited other imaging modalities, such as fMRI [51]. Simulation-based approaches, however, often suffer from being either too simple or too complex. While simple simulations are computationally efficient, they often only create realizations of the model being tested rather than the actual underlying phenomenon [52, 53]. Complex simulations instead capture too much detail and are severely limited computationally, requiring high-performance computing to simulate more than small volumes with a handful of neurons [54, 55]. For these reasons, existing methods do not provide plausible and computationally efficient simulations useful for large-scale functional imaging.

To assess TPM methods with realistic and computationally efficient simulations, we present the Neural Anatomy and Optical Microscopy (NAOMi) simulator. Our framework leverages simple, but flexible, models of neural tissue to efficiently create large volumes with thousands of neurons on standard workstations (Fig. 1). Arbitrary patterns of spiking activity can be generated for this population, which our framework then transforms into realistic fluorescence traces separately for somas as well as processes. A light model approximates laser propagation and scattering throughout different locations of the simulated tissue. These components are combined in a simulated scanning procedure that incorporates important imaging effects, such as sample motion. We describe the simulation model, which has a publicly available software implementation^1^, and provide parameters for simulating two-photon GCaMP [46] recordings in layer 2/3 of mouse visual cortex. We use these simulated datasets to evaluate several automated calcium imaging demixing algorithms. Finally, we generate several more datasets to compare the performance of standard and specialized TPM experimental setups under a variety of sample conditions.

**Figure 1:**
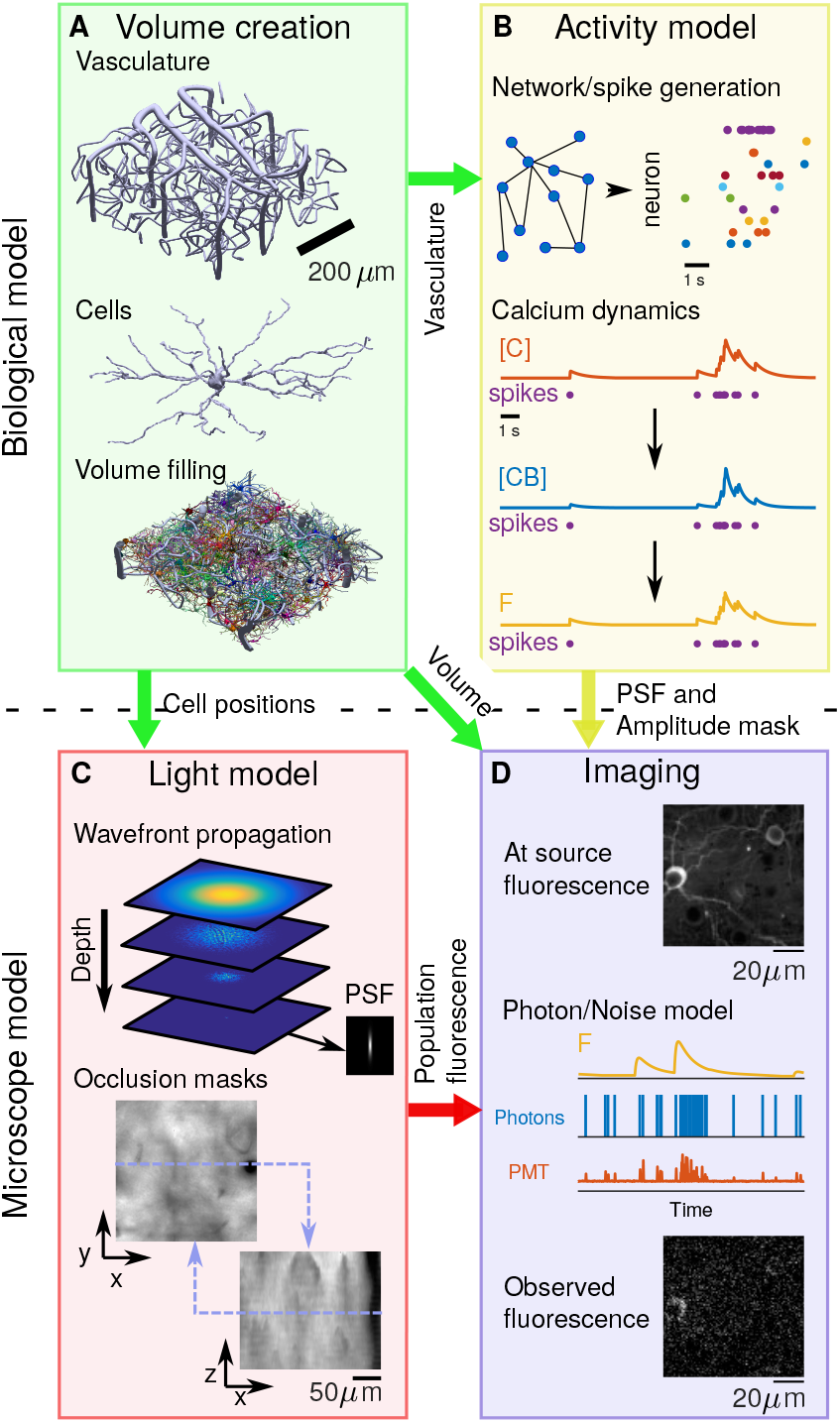
Block diagram of NAOMi simulator. A: Neural volume generation process. Vasculature is generated throughout the volume, followed by cell bodies and finally dendrites and axons are grown. B: Network activity generation. The spiking activity for each neuron is simulated and converted into calcium ([C]), bound calcium ([CB]), and fluorescence (F) for a chosen indicator. C: Light propagation model. An optical wavefront corresponding to particular microscope optics is propagated through a simulated scattering volume, generating a scattered point-spread function and relative intensity masks. D: Scanning and image formation. The volume, modulated by the simulated activity, is scanned using the output of the light model with motion and noise sources from a model of the light collection, amplification, and digitization process.

## 2 Results

### 2.1 Simulation design

Generating realistic imaging data useful for honest assessment of a spectrum of techniques hinges on accurate, efficient simulations of anatomical volumes at the scale of optical imaging (Fig. 1A). Our anatomical simulation starts by constructing a scaffolding of vasculature with three parts: surface vessels, diving vessels, and capillaries [56] (Tab. 1, Sup. Fig. 1, 2, 3, See Methods). Next neurons are placed throughout the volume, first placing somas, and then growing dendrites and axons [57] from the cell bodies (Sup. Fig. 4, 5, See Methods). Statistical models of neurons and process paths ensure variation in cell shapes and were tuned using morphological data from electron microscopy (EM) data [58] and optical microscopy [59–61] (Tab. 1, See Methods).

**Table 1:** Parameters used for *in-silico* simulation of neural activity in layer II/III of mouse primary visual area V1. Values for each parameter were either directly found in the literature or estimated from published data (entries with a †). The third column indicates whether these parameters were set directly in NAOMi, or were fit indirectly by setting other simulation parameters. In the latter cases, the measured values from a simulated NAOMi volume are shown for comparison, indicating that the simulated anatomy matches measured anatomical statistics.

| Parameter | Lit. Val. | Fit type | NAOMi Val. | Unit | Refs |
| --- | --- | --- | --- | --- | --- |
| Anatomical parameters: |  |  |  |  |  |
| Neural density | 9.20E+04 | Direct | – | mm<sup>-3</sup> | [58] |
| Fraction vasculature | 0.01-0.04 | Indirect | 0.032 | – | [58, 74, 75] |
| Fraction cell bodies | 0.12 | Indirect | 0.135 | – | [58] |
| Fraction neuropil | 0.84 | Indirect | 0.833 | – | [58] |
| Fraction dendrites* | 0.294 | Indirect | 0.223 | – | [58] |
| Fraction other (fluorescing)* | 0.401 | Indirect | 0.33 | – | [58] |
| Fraction other (not fluorescing)* | 0.293 | Indirect | 0.28 | – | [58, 76] |
| Vessel radius (capillary) | 2.00E+00 | Direct | – | μm | [56] |
| Vessel radius (penetrating) | 10 (9,11) | Direct | – | μm | [56] |
| Vascular density | 1-3 | Indirect | 2 | % | [56, 58, 74, 75] |
| Penetrating vessel density | 30<sup>†</sup> | Direct | – | mm<sup>-2</sup> | [56] |
| Somatic volume* | 1.80E+03<sup>†</sup> | Indirect | 1.80E+03 | μm<sup>3</sup> | [77] |
| Nuclear volume* | 800<sup>†</sup> | Indirect | 800 | μm<sup>3</sup> | [77] |
| Cytoplasm volume* | 1000<sup>†</sup> | Indirect | 1000 | μm<sup>3</sup> | [77] |
| Basal dendrite diameter | 0.7 | Direct | – | μm | [78] |
| Basal dendrite length | 100-160<sup>†</sup> | Indirect | 105 | μm | [59–61] |
| Apical dendrite diameter | 1-2 | Direct | – | μm | [79] |
| Axonal diameter | 0.3 | Direct | – | μm | [58] |
| Fluorescence parameters: |  |  |  |  |  |
| GCaMP6f binding affinity $K_d$ | 290 | Direct | – | nMol | [70] |
| Baseline Ca<sup>2+</sup> concentration | 50.00 | Direct | – | nMol | [68] |
| Ca<sup>2+</sup> binding ratio $k_s$ | 100,110 | Direct | 110 | AU | [67, 80–82] |
| Ca<sup>2+</sup> diffusion constant $\gamma$ | 1800 | Data fit | 292.3 | s<sup>-1</sup> | [67, 80, 83] |
| Ca<sup>2+</sup> axon diffusion constant $\gamma$ | 2800 | Direct | – | s<sup>-1</sup> | [67, 80, 81] |
| GCaMP6f Hill eqn. exponent $n_h$ | 2.7 | Direct | – | AU | [70] |
| GCaMP6f Hill eqn. amplitude | 25.2 | Direct | – | F | [70] |
| Indicator concentration | 10-200 | Direct | 10 | μM | [68, 84, 85] |
| *Adjusted for shrinkage: 31% [86], <sup>†</sup>Value estimated from data in the literature |  |  |  |  |  |

The next step in simulating TPM data is to augment each generated neuron with realistic fluorescence activity (Fig. 1B). Spiking activity for each neuron is either pre-defined or is generated using models that output correlated, bursting population activity based on models of neural connectivity [62–66] (See Methods). Next, the known non-linear calcium decay process simulates the dynamic concentration of calcium ions [67, 68]. This two compartment model describes separate dynamics for the cell bodies and the neurites, both driven by the same spike trains. As in related work, a protein-specific double exponential model modulates the calcium concentrations to create bound calcium concentrations with appropriate onset and offset time-constants [68, 69]. Finally, the bound calcium concentrations are converted to fluorescence values using the Hill-equation fit to fluorescence measurements [70, 71] (Tab. 1, Sup. Fig. 8, See Methods).

The next step in the simulation is to estimate the optical properties of the specified microscope configuration within the generated tissue (Fig. 1C). The scattering nature of brain tissue substantially affects light propagation through it, resulting in an abberated point-spread function (PSF) and decreased optical performance. We approximate these effects by performing wavefront propagation of a specified beam shape (i.e. Gaussian or Bessel beams [72, 73]) through a generated volume of refractive index shifts (Sup. Fig. 10, See Methods), generating simulated PSFs across the volume. The weight of the simulated PSF across different locations of the simulated volume forms an occlusion mask, representing inhomogeneity of optical performance across the sample. This occlusion mask is also modulated by an estimate of the absorption of emitted light through blood vessels and the neural volume as a function of position.

The final scanning module combines the outputs of the anatomical, light, and activity modules to produce images on a frame-by-frame basis (Fig. 1D). The fluorescence activity and occlusion mask modulate the anatomical volume, which is convolved with the simulated PSF to produce raw, noiseless, illumination images. Sub-pixel line-by-line offsets, representing brain motion, are applied prior to spatially resampling to the desired image resolution and applying the measurement noise model. Measurement noise is simulated by per-pixel Poisson sampling of photons counts at the photo-multiplier tube (PMT) and converting these counts into electrical measurements via PMT photon and electronics amplification distributions (Sup. Fig. 10, See Methods). This process includes bleed-through across pixels from the amplifier’s temporal response kernel (Sup. Fig. 12, See Methods). This procedure is performed independently for each frame, and captures the complex, non-Gaussian noise profile inherent in TPM data.

### 2.2 Comparison of simulated data to real data

We next evaluate the overall simulation output against recordings from mouse V1 (Fig. 2). We generated a 500 *μ*m x 500 *μ*m x 100 *μ*m volume and scanned it with a 0.6-NA Gaussian point-spread function over 20,000 frames at 30Hz sampling and 40mW laser power, comparable to parameters in a recorded dataset obtained from mice expressing GCaMP6f being exposed to a set of visual stimuli (See Methods).

**Figure 2:**
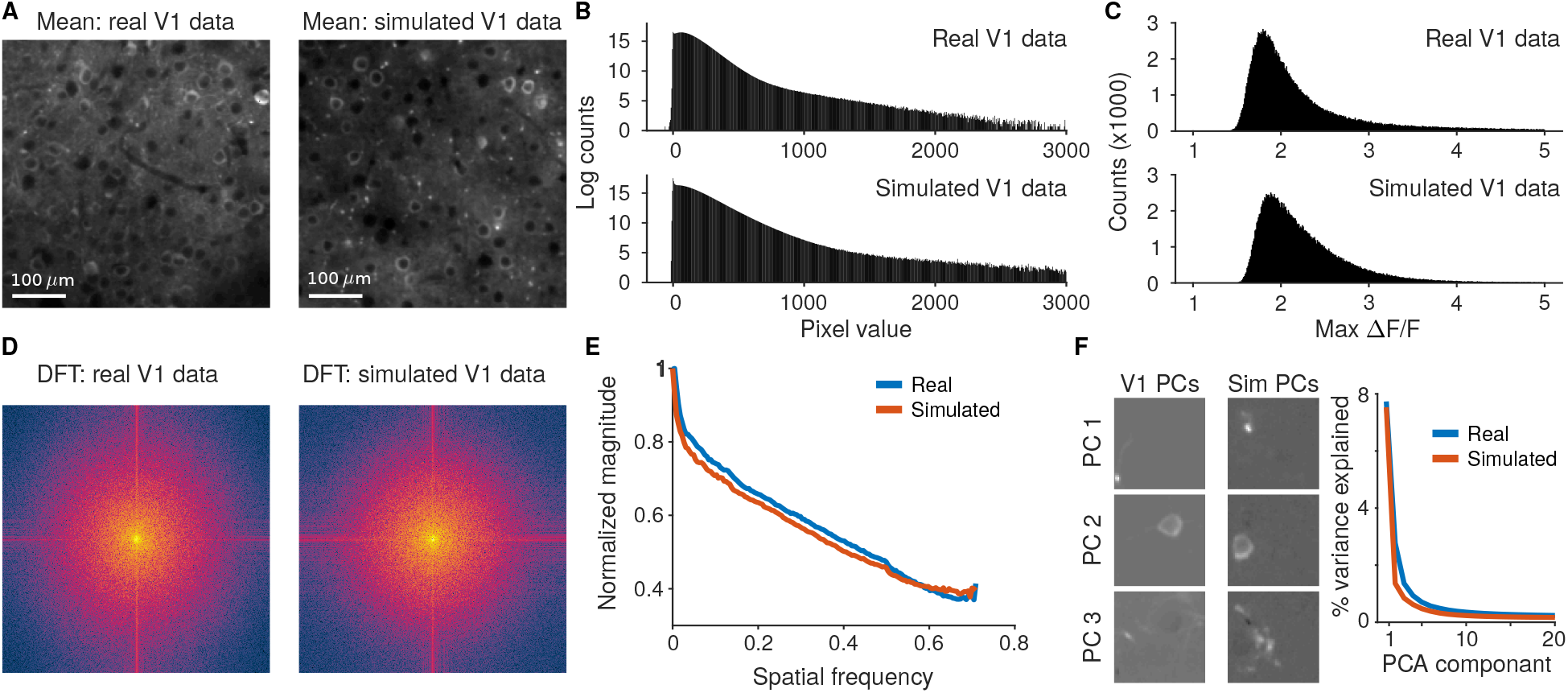
Comparison of simulated data to recordings of mouse V1 L2/3 using GCaMP6f. A: The mean image for mouse V1 recordings and simulated data. B: Pixel value distributions across the full videos display bimodal peaks and a right log-linear tail. C: Distribution of the maximum ΔF/F values across all pixels in the FOV match between the simulated and real V1 data. D: The spatial frequency content in the mean simulated image captures the qualities of the real data. Both the spread of frequencies and the tendency for high-frequency components in the fast- and slow-scan directions that result from line-by-line motion and pixel bleed-through are captured. E: The overall contributions at different spatial frequencies to the mean activity matches between the recording and simulation. F: Principal component decompositions for both the real and simulated data exhibit similar decays in the variance explained per component. The resulting spatial principal components are qualitatively similar.

The simulated videos and recorded videos visually share many of the same features (Sup. Video 1), including bright, sparse transients of fluorescence across the whole image. The overall mean images (average of frames across time; Fig. 2A) both show distinct cell bodies along with muted processes that have their intensity modulated by scattering from blood vessels and other tissue elements. Histograms of video pixel values (Fig. 2A,B) feature heavy right tails corresponding to neural activity and also peaks at zero corresponding to zero-photon pixels.

The neural activity distribution for individual pixels is explored by comparing the relative strength of firing activity across the field of view (FOV). The distribution of maximum activity (maximum ΔF/F over 20,000 frames) for all pixels (Fig. 2C) for each of the two videos is peaked at 2 with a slight heavy right tail, which corresponds with neurons that fired large transients within the videos. Other statistics, such as the distribution of values in the mean image, the standard deviations over all pixels, and measures of activity such as the ratios of maximum to median fluorescence values also match well (Sup. Fig. 14A-C).

The global frequency content of the two videos is estimated with the 2D discrete Fourier transforms of the mean images (Fig. 2D). The Fourier transforms of both videos depict very similar features, such as increased frequency content along the fast- and slow-scanning axes resulting from the sequential pixel bleed-through and residual line-by-line motion artifacts. Additionally, the frequency fall-off (Fig. 2E) for both the real and simulated data display the same decay. Finally, observing the effective dimensionality of both videos via Principal Component Analysis (PCA; Fig. 2F) finds both qualitative similarities between the spatial principal components (PCs) and quantitative similarities between the distribution of variance explained for the leading PCs on small patches of videos.

### 2.3 Evaluation of automated segmentation

We start evaluating TPM techniques using NAOMi by analyzing the performance of automated demixing algorithms, leveraging the ground truth information available. We applied three common algorithms — PCA/ICA [29], constrained non-negative matrix factorization (CNMF) [38], and Suite2p [18] — to 20,000 frames simulated from a 500 *μm* x 500 *μ*m x 100 *μ*m volume with 1 *μ*m sampling at 30Hz scanning using a 0.6-NA Gaussian excitation numerical aperture (NA) at 40 mW average power. The ground truth consisted of the spatial profiles of each individual neuron and component within the volume and their individual fluorescence traces.

Each algorithm returned a set of demixed time traces and corresponding spatial profiles (Tab. 6–8, Sup. Fig. 16–19). Overall CNMF, Suite2p and PCA/ICA isolated 1091, 661, and 265 components, respectively, out of a total of 8,117 possible fluorescing components. Comparisons to the ground truth traces, based on a combined Pearson’s correlation cut-off of 0.1 on the time-traces and a 50% pixel overlap, revealed which components represented actual cells in the volume (Tab. 6). We considered a pairing to be a “strong pairing” if the correlation exceeded 0.5 (Fig. 3A,B). These correlation values account for all aspects of how well the estimated traces match the true time-courses, including missed transients and false transients from other components (e.g., neuropil; Fig. 3C).

**Figure 3:**
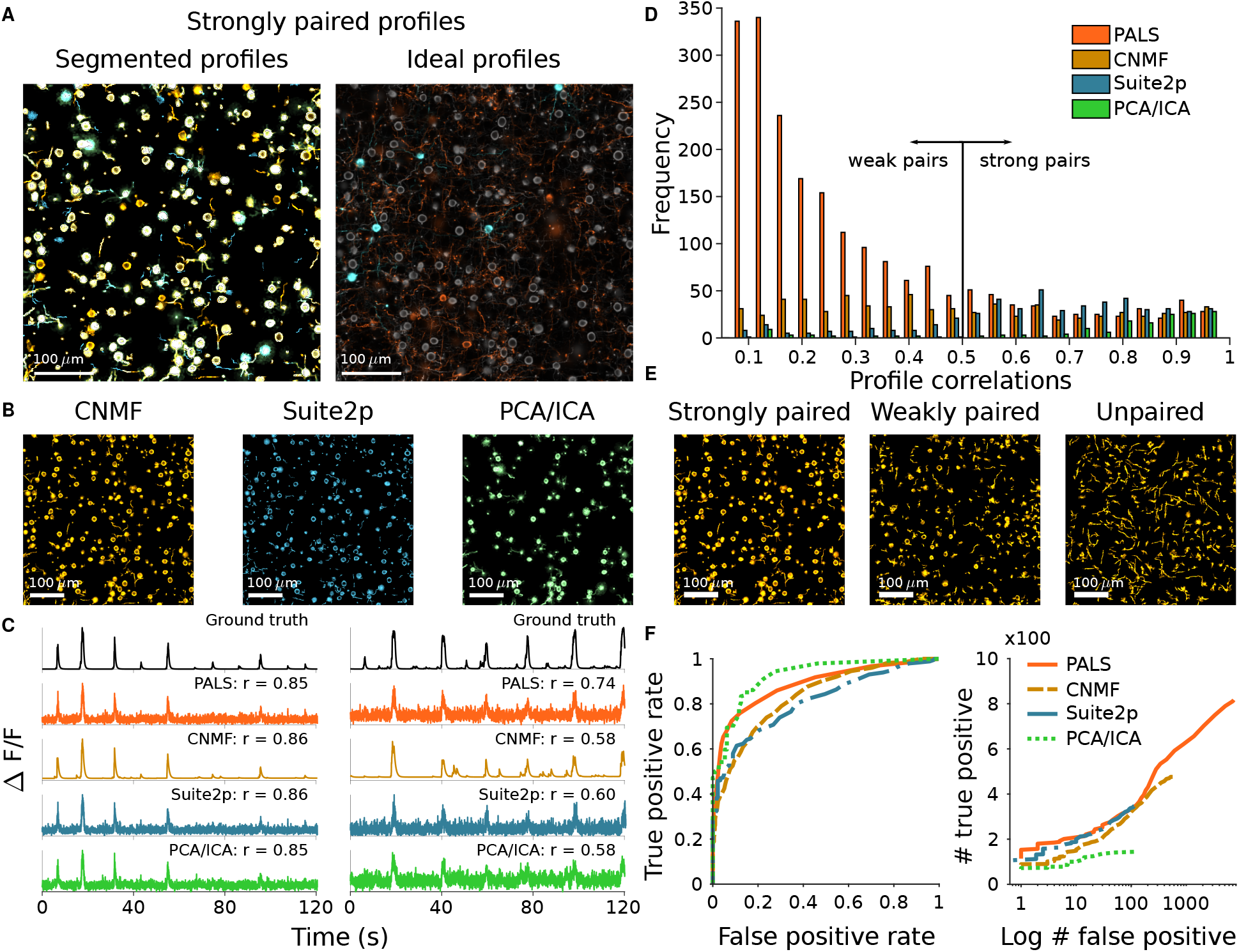
Comparison of popular calcium imaging segmentation algorithms using synthetic data generated using NAOMI. A: Spatial profiles from CNMF (red), Suite2p (blue), and PCA/ICA (green) overlaid with ideal profiles (yellow) calculated from the simulation volume. B: Strongly paired (r >= 0.5) spatial profiles from each algorithm displayed separately. C: Example timecourses estimated by each of the segmentation algorithms as compared to the ideal profile assisted least-squares (PALS) estimated timecourse and the ground truth timecourse. D: Histogram of correlation values of estimated timecourses to the ground truth timecourses for each match spatial profile. E: Spatial profiles from CNMF separated into strongly paired (r>=0.5), weakly paired (r<=0.5) or unpaired. F: ROC curves for strongly paired (r>=0.5) spatial profiles sorted by their peak fluorescence and profile weight.

Of the paired profiles, some were doubled, i.e. multiple algorithmically discovered profiles matched to different portions of the same simulated cell (Sup. Fig. 23, Tab. 6). Accounting for doubling, CNMF, Suite2p, and PCA/ICA found 303, 292, and 137 unique cells at the > 0.5 correlation level. Interestingly, while CNMF finds the most distinct components (i.e., before accounting for cells found with multiplicity), it only finds approximately the same number of unique cells as Suite2p, and both have a lower rate of found true cells than PCA/ICA (Fig. 3D). Furthermore, comparisons of individual cells found (Tab. 7, 8) show that different methods find non-overlapping sets of cells (Sup. Fig. 21). For example, CNMF and Suite2p only agree on 273 of the ≈ 300 cells (Fig. 3A, Tab. 7).

While these figures may seem small compared to the 8,117 total sources, not all fluorescence sources are visible above the noise level. To explore this effect with NAOMi, we computed auxiliary time-traces from the raw, noisy video using the “ideal” ground-truth spatial profiles to obtain the profile-aware least-squares (PALS) time trace estimates (See Methods). Due to the video signal-to-noise ratio (SNR), these estimates yielded only 415 timecourses accurately matched at the > 0.5 correlation level (Fig. 3D, Tab. 8) indicating that the gap induced by simultaneous estimation of spatial profiles is not overly large. In fact, the inherent denoising in some algorithms allows some cells’ time courses to be estimated with even higher fidelity than the traces derived from the ideal spatial profiles (e.g. CNMF identified 8 cells at the *r* > 0.5 level that the ideal profiles produced lower correlation values for; Tab. 8).

One challenge in interpreting the results of automated demixing is that, sans ground-truth, it is difficult to determine if a source is a true cell or an artifact. Instead, sorting components based on metrics such as overall fluorescence levels can be used. Varying one such criterion — a threshold on the maximum fluorescence — to classify true and artifact sources results in receiver-operator characteristic (ROC) curves that compare the number of strongly paired components kept (true positives), to the number of weakly paired or unpaired components kept (false positives). These curves show that while PCA/ICA obtained the fewest components overall, CNMF and Suite2p found bright artifacts at much higher rates (Fig. 3E, Sup. Fig. 24).

One benefit of the NAOMi simulator is that we can easily explore how optical parameters effect algorithmic performance. We replicated the above analysis with a 2x increase in laser power (80 mW), keeping the volume and neural activity constant. Ideally this power boost would illuminate additional cells, as weaker and more sparsely firing cells would be more distinguishable. We found that all algorithms returned more unique components at the *r* > 0.5 level, with 424 for CNMF, 358 with Suite2p and 264 for PCA/ICA: a 39.93%, 22.6%, and 92.7% improvement (Sup. Tab. 9), respectively. Interestingly, a similar increase in components that did not pair well to real cells meant that there was negligible improvement in rate of found cells, and some ROC curves reduced in area, indicating that fluorescence magnitudes became less sufficient to differentiate true cells from artifacts (Sup. Fig. 24.

We note that in addition to laser power, other factors such as the sampling resolution, numerical aperture and neuropil strength also influence the ability to detect neural activity. NAOMi enables exploration of all these aspects. For example we find 1) a sharp cut-off in the ability to accurately detect components when sampling at intervals larger than 3μ m (Sup. Fig. 25) 2) an improvement in the ability to detect smaller burst sizes in the ΔF/F values with neuropil correction as in Suite2P [18] (Sup. Fig. 26), and 3) a steady decay in signal strength per component as a function of NA, reaching a critical reduction of signal at NA ≈ 4 (Sup. Fig. 27).

### 2.4 Evaluation of TPM optical configurations

The ability to modify optical parameters and sample expression patterns allowed for direct assessment of the trade-offs between microscope configurations across sample conditions. We applied this new mode of assessment to perform three head-to-head comparisons: 1) imaging of sparsely labeled tissue using Bessel [8] vs. high-NA Gaussian beams (Fig. 4A-C), 2) imaging of nuclear labeled tissue using high-NA vs. low-NA (axially extended) Gaussian beams (Fig. 4D-F), and 3) volumetric imaging of densely labeled tissue using multiplane Gaussian [87] vs. temporally-focused beams [88] (Fig. 4G-I).

**Figure 4:**
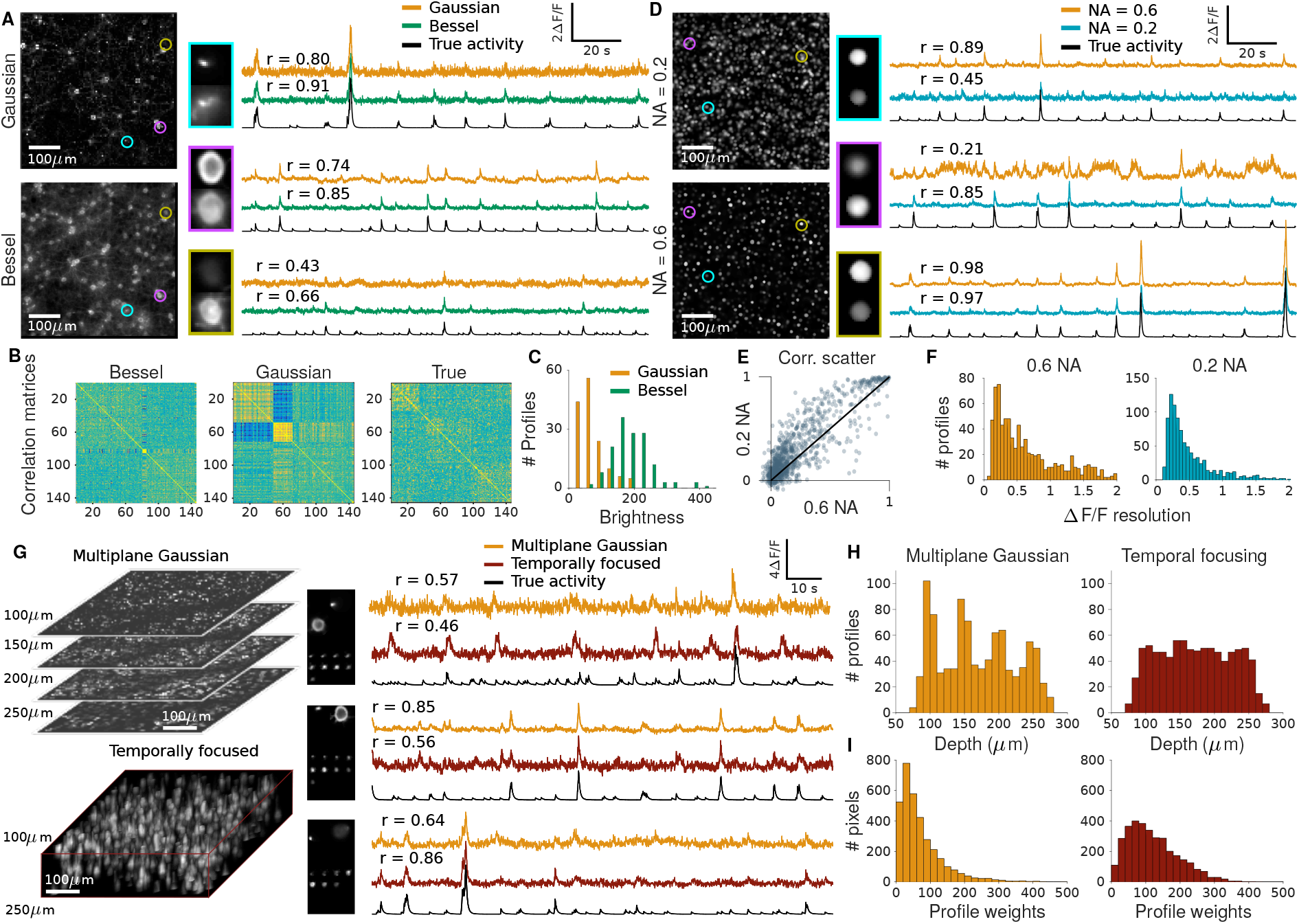
Comparison of specialized imaging modalities to standard high NA Gaussian TPM for a sparsely labeled volume (A-D), nuclear labeled volume (E-H), and volumetric imaging (I-L). A: Mean image (left) and example time traces (right) of a sparsely labeled volume with Bessel beam (top) and high NA Gaussian (bottom) illumination. B: Correlation matrices of extracted time traces for each method sorted by clustering into 3 groups using k-means. C: Histograms of spatial profile weights of cells in the volume using Gaussian and Bessel PSFs. D: Mean image (left) and example time traces (right) of a nuclear labeled volume with low NA Gaussian (top) and high NA Gaussian (bottom) illumination. E: Scatterplot of correlation values for low and high NA extracted timecourses against the true timecourse. F: Estimated Δ*F*/*F* resolution of cells based on their spatial profile and the mean image. G: Mean image (left) and example time traces (right) of volumetric TPM with 4 high NA Gaussian planes (top) and 16 temporally focused planes (bottom) illumination. H: Histogram of axial positions of highly correlated (r > 0.5) cells using a scanned high NA Gaussian or temporally focused illumination. I: Histograms of profile weights using high NA Gaussian or temporally focused illumination.

In our first comparison we simulated TPM recordings of a sparsely labeled (10% neurons expressing GCaMP6f) 500 *μ*m x 500 μm FOV of mouse cortex using both conventional high-resolution TPM with a Gaussian PSF (0.6 NA) and extended depth-of-field TPM using a Bessel beam (0.4 NA, 60 *μ*m long) [8]. As shown previously, Bessel beam imaging resulted in excitation throughout the whole volume, resulting in more uniform excitation of neurons (Fig. 4C) and more neurons recorded with high signal fidelity. Another consequence of the uniform excitation observed using Bessel beams is an increased robustness to axial motion. To test this, we clustered the extracted time-traces from Bessel and Gaussian imaging into three groups (Fig. 4B). The sorted correlation matrices showed a strong motion-induced artifact only for Gaussian imaging, suggesting a reduced influence of motion artifacts on time-trace estimation using Bessel beams.

Our second comparison tested the performance of nuclear labeled TPM, which has been extensively used for larval zebrafish and *C. elegans* TPM [89, 90] but not in mouse brain imaging, in a 500 *μ*m x 500 *μ*m FOV of mouse cortex. We imaged this simulated tissue using Gaussian beams of two excitation NAs: 0.2 and 0.6 (Fig. 4D). Surprisingly, simulations of signal levels across a variety imaging NA values showed constant overall integrated value (i.e., signal level; Sup. Fig. 27), suggesting little loss of power efficiency by reducing the excitation NA. Similar to the extended depth-of-field Bessel beam, the 0.2 NA excitation resulted in sampling more neurons from the volume and improved overall imaging performance (Fig. 4E,F). As the imaging volume is mostly non-fluorescing with nuclear-labeling, distinguishing individual cells is straightforward even with a extended depth-of-field. Because nuclei are several times larger than the lateral resolution of the 0.2 NA excitation beam, there are fewer advantages to switching to a Bessel beam setup and low NA imaging is more power efficient.

Last, we compared multiplane TPM [87] to scanned temporal focused (s-TeFo) TPM [88] in a 500*μ*m x 500 *μ*m x 200 *μ*m volume of the mouse cortex layers 2-4 (Fig. 4G). For multiplane Gaussian TPM, we scanned four planes, separated by 50*μ*m at 1 *μ*m lateral sampling, at 10Hz and for s-TeFo TPM, we scanned 16 planes, separated by 10*μ*m at 4*μ*m lateral sampling, at 10Hz. Given a similar amount of excitation within the volume, both techniques performed comparably in extracting fluorescence traces with high fidelity. The axial distribution of highly correlated (r > 0.5) cells in the multiplane TPM is multimodal, set at the focal positions the four planes, while the s-TeFo distribution is much more uniform (Fig. 4H), as is reflected in the profile weight histograms (Fig. 4I). These simulations suggest that while s-TeFo as a technique may not drastically increase the number of recorded neurons, the more consistent spatio-temporal sampling of neurons may decrease sampling bias and provide robustness to motion-induced artifacts and signal crosstalk.

## 3 Discussion

We present here the NAOMi simulation framework that generates detailed TPM data. This framework yields new insights into TPM technology and enables the testing and validation of existing and novel TPM methods, allowing for assessment and methodological optimization not currently possible. Toward this end, we have developed the ability to generate realistic synthetic neural volumes, transient calcium activity, and two-photon calcium imaging videos. We further increase the simulation efficiency by simplifying statistical models of the processes involved, reducing the computational burden and making this tool more broadly applicable.

We have demonstrated two important use cases of NAOMi: assessment of calcium imaging demixing methods and comparing optical configurations across imaging conditions. In both cases, useful large-scale ground truth is difficult to obtain experimentally. NAOMi leverages the accumulated knowledge of neuroanatomy, optical physics, and neuroscience to bypass these difficulties through simulation.

To assess demixing algorithms, we generated a calcium imaging dataset typical of TPM in mouse neocortex and analyzed the performance of several popular methods: CNMF [52], Suite2p [34], and PCA/ICA [29]. The available input cell shapes and timecourses enabled direct comparisons of the algorithm decompositions to ground truth, which revealed that methods with the highest number of found neurons also had much higher instances of artifactual components. This indicates the potential for many false positives in automated demixing (Sup. Fig. 20, Tab. 6). Using NAOMi data as a testing ground, more advanced detection metrics can be developed, to find better ways to filter out true cells from the full set of returned components.

Using NAOMi, we explored particular combinations of optics and samples *in silico* and illustrated the advantages three specialized techniques could have while imaging particular sample conditions. We rapidly assessed whether or not a particular technique is appropriate for the sample and optimized imaging parameters to maximize the quantity and quality of the data we collect. Additionally, we explored the interaction of different components of *in vivo* calcium imaging and their effects on performance. For instance, we quantified the effects that axial brain motion can have on estimated cell-cell correlation as a function of both sample type and optical configuration.

Despite these results, there are several aspects of the simulation that were simplified for speed or simplicity. While we provide a rudimentary spike-train simulation to drive the fluorescence models, we note that more realistic firing patterns for neurons will help improve overall realism. A number of complex packages simulate such activity for neural populations (e.g. [91]). In these cases, rather than replicate these methods, we encourage their use and integration into the NAOMi framework.

In the development of NAOMi we aimed to create a tool accessible to the community at large that is easily expandable in its scope and abilities. To this end, our software was designed to be modular so that as better anatomical models, optical descriptions of tissue, and TPM statistics become available, they can be easily incorporated into the existing framework. Additionally, single modules can be modified in order to simulate different setups. For example, changing the anatomical structure (e.g., blood-vessel size and cell body statistics) can allow for bench-marking imaging techniques in rats, rather than mice. These changes and extensions will allow NAOMi to be a useful tool for a wide variety of applications for experimentalists and methods developers.

Work in other fields has shown the great utility of developing strong simulation-based models of experimental data (see [92] and references therein). NAOMi is a tool that fills part of this gap for neuroscience data. As neuroscience continues maturing, better models of data must be developed, especially for data that is as diverse and complex as two-photon calcium imaging. This and other work will allow researchers to not only more quantitatively judge the quality of their data, but also make better predictions on the data they will need for their experiments.

## 4 Materials and Methods

Our TPM simulator is designed to permit testing of many different aspects of the calcium imaging process. To achieve this flexibility, our simulator is divided into five distinct modules, each focused on a portion of either the tissue or scanning simulation (Fig. 1). The five modules are: 1) the neuron module responsible for generating single neurons, 2) the volume module responsible for assembling the neurons into a tissue volume that includes neuropil and vasculature, 3) the activity module that generates the temporal calcium traces for each neuron and the neuropil, 4) the optics module that simulates the point-spread function and occlusion due to the optical mask, and 5) the imaging module that simulates the TPM noise model and object motion.

### 4.1 Neuron model

The first module creates simple, yet plausible models of neurons that can be placed throughout a volume and scanned in simulation. We model the neural shape via a probability distribution over smooth deformation of a sphere, followed by a nonlinearity. This model allows for fast sampling of unique neurons, meaning that each simulated volume will contain a completely new set of neurons. Additionally, we provide for each neuron a nucleus modeled as a shrunken and smoothed version of the soma shape. This model captures the relationship observed in detailed field emission scanning electron microscopes (FESEM) [93]. Finally, we simulate for each neuron a number of dendrites, one of which is created thicker and at a downward orientation, as to model the apical dendrites.

The model of the smoothly deformed cell body is an isotropic Gaussian process [94] defined over a sphere. To sample from this distribution and create the cell body, we sample uniformly over a sphere [95], sample *i.i.d*. a Gaussian random variable for each point, and smooth the points according to the process covariance. Denoting the sample points ***p***_*i*_ ∈ ℝ^3^, the height (distance from center of the sphere) can be sampled from

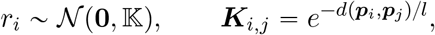

where *l* is the length-scale that controls the smoothness of the cell body, and *d*(·, ·) is the geodesic distance between any two points. For the unit sphere (radius one), this distance is the arc length along the great circle connecting the two points

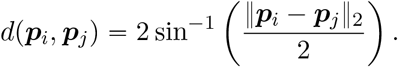

When unconstrained, the radial height of this function can, at times, exceed the maximum and minimum realistic deformations *r*_max_ = max_*i*_|*r_i_*| and *r*_min_ = min_*i*_|*r_i_*|. We thus rescale the radii values as

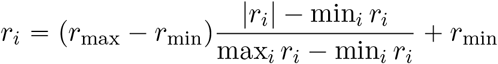

The resulting points *r_i_**p**_i_* form the points for a mesh that define the interior of the cell body. To account for the non-spherical shape found in pyramidal neurons, we can modify the radii values by making the base radius at each point dependent on a function of its location on the sphere. Specifically, we use the equation for a tear-drop that is defined parametrically by the azimuth and elevation angles *ϕ*, and *θ* as

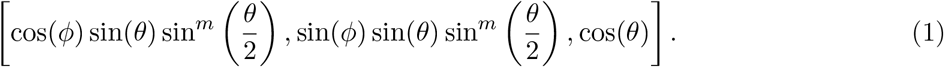

The final step in creating the cell body is to create the nucleus, which is accomplished by shrinking and smoothing the cell wall shape as defined by *r_i_**p**_i_* as

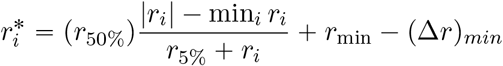

Dendrites are added to each neuron via a stochastic growing process [57]. The process generates start and end points for each dendrite, and iteratively grows the dendrite through the volume while avoiding any obstacles (i.e. other cell’s somas, dendrites, or blood vessels). Apical and basal dendrite endpoints are separately set within the volume and the grown dendrites are dilated to widths consistent with measured anatomy (Tab. 1, Sup. Fig. 7).

### 4.2 Volume generation

To create the tissue volume, we initialize an empty volume and begin by placing blood vessels throughout the volume. For computational feasibility, the volume is modeled as a 3-D grid of points with sub-micron sampling (we typically use 0.5*μ* m distances). The blood vessels are grown in three parts: surface vasculature, diving arterioles, and capillaries. Surface vasculature is grown by connecting nodes randomly placed upon the surface of the volume. The connected paths are smoothly varied and dilated. Diving arterioles are set at endpoints of surface vasculature and connected to the bottom of the simulated volume. Capillaries are connected to the diving arterioles and pseudo-randomly placed within the volume in a space-filling fashion. Vessel diameters, concentration, branching frequency, and orientation were compared and fit to two-photon microscopy data of mouse vasculature (Schaffer-Nishimura lab, unpublished data).

Once initialized, the volume is then filled with the neuron somas. We sequentially place the neurons randomly throughout the empty space in the volume, with a minimum distance that allows cell bodies, but not nuclei, to overlap. The random placement can be modified to encourage neurons to be more spread out, or more cluttered. When an overlap occurs, the overlapping region is given to the latest cell to be placed. This allows our volume to contain touching cell bodies. Once all the cell somas are placed, dendrites are grown for each neuron sequentially, such as to avoid location conflicts with other cells. Apical dendrites are grown in the same fashion, only thicker, axially oriented, and having fewer transversal deviations. Separate apical dendrites corresponding to neurons in deeper cell layers are grown in a similar fashion from the bottom of the volume to the top.

As a final step, axons fill up the remaining empty space, up to the typical 0.7 fill fraction of layer 2/3 in mouse V1. The same dendrite growing algorithm [57] is used to create millions of short axon segments throughout the entire volume. To obtain the global correlated background components, we overlay 3D Gaussian processes that indicate the strength of each component throughout the volume with the set of axon locations. Specifically the value of the *k^th^* background component

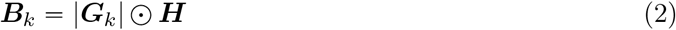

where ⊙ represents element-wise multiplication between the 3-tensors ***H***, which contains a one where a neuropil process is present and zero otherwise, and the absolute value of ***G***_*k*_, a GP with mean zero and Gaussian radial-basis function covariance kernel where the covariance between any two points at locations ***p***_*i*_ and ***p***_*j*_ is

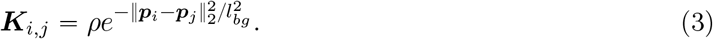

### 4.3 Time-trace generation

To simulate temporal activity, we provide a number of options to generate time-traces for each neuron. We provide both statistical models that generate stereotypical activity as well as more detailed [Ca^2+^] dynamics model. The statistical model provides a simple way to input basic behaviors of various fluorescent proteins (i.e. rise-time and decay). The [Ca^2+^] dynamics model simulates the molecular kinetics over time, and provides a way to test the time-trace assumptions made in calcium imaging analysis algorithms.

#### 4.3.1 Spike-time generation

We provide two methods to generate spike trains to drive the fluorescence activity simulation. The first method creates independent activity for each neuron, including bursting behavior. The second model simulates a Hawkes process which accounts both for self-excitation, driving busting behavior, as well as inter-neuron spiking correlations [96].

To generate independent spike trains, we model each neuron as a bursting neuron, where bursts occur at independent, exponential intervals

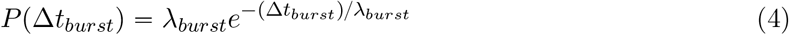

for Δ*t_burst_* > 0. The rate of bursting λ_*burst*_ is chosen differently for each neuron. The rates can be given to the simulator, or the simulator can automatically draw burst rates from a Gamma distribution with a provided mean rate and parameter *α* = 1. For each burst, the number of spikes are chosen as

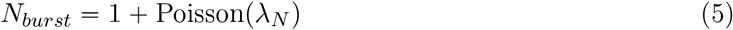

where the parameter λ_*N*_ controls the length of the bursting. The inter-spike times between spikes in a burst were modeled as uniformly random between 5 ms and 7 ms. Alternative distributions of spiking activities can easily be implemented by passing different λ_*burst*_, *α*, or λ_*N*_ values to the simulator, or by direct modification to the code to implement different distributions that better reflect activity in other cortical areas.

For the Hawkes model simulation, we first generate a connectivity matrix that encodes how each neuron’s firing excites other neurons. We model this connectivity with a Watts-Strogatz small-world network model [62]. To correlate the processes to the network activity, we allow for all neurons to influence the background processes, while not allowing many return connections. To stabilize the point-process, we normalize the resulting connectivity matrix to have maximum eigenvalue magnitude of 0.98. We then run the Hawkes process using Lewis’ method [97], with an exponential distribution over the neuron’s base firing rates and a higher base firing rate for the background components. Finally, we bin the resulting continuous-time spike events into 1 ms bins to create the discretized spikes that are then fed into the calcium dynamics simulation.

#### 4.3.2 AR-*p* dynamics

For each cell, we generate a baseline fluorescence, *β_i_* = |1 + *z*| where 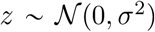 is a Gaussian random variable. The variance *σ*^2^ controls the distribution of baselines, and we set a default value to *σ*^2^ = 0.04. The next step is to simulate the spike or “event” times for each neuron. As most neurons are sparsely active, we draw the firing rate of each neuron as λ_*i*_ ~ Gamma(*a, θ*). The parameter *θ* gives the average inter-spike distance in time and should be set according to the temporal sampling rate set in the simulation. The parameter *a* is the shape parameter and modulates the distribution of the firing rates. We find that *a* = 1 (where the Gamma distribution collapses to an exponential distribution) yields realistic distributions of neuron activity levels. The actual event times are then sampled for each neuron according to a Poisson process with rate λ_*i*_. To model the different calcium levels at each event (e.g. due to multiple spikes or to adaptation [98]), we sample the overall concentration as coming from a unit log-normal distribution (i.e. an exponentiated Normal distribution 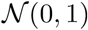).

Once the spike times are obtained, an auto-regressive model with *p* degrees of freedom (AR-*p*) is used to simulate the calcium and fluorescence impulse response. As a difference equation, AR-*p* models can be written as

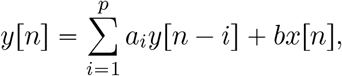

where the *a_i_*’s are the AR coefficients and *b* is a scalar multiple of the input. The impulse response can be obtained by solving the inverse Laplace transform

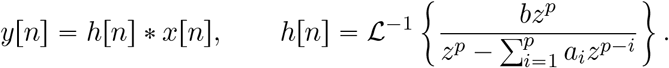

Standard linear systems theory shows that *h*[*n*] will be composed of the exponentiated roots of the characteristic polynomial 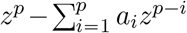, and therefore will be an exponentially decaying function. Higher order polynomials can result in a rise time as well. For this work we find that an AR-2 model (*p* = 2) sufficiently models the rise and fall of observed GCAMP responses. The filter ***h*** is convolved with the spike-time vector to create the temporal activity per neuron.

#### 4.3.3 [Ca^2+^] dynamics

The fluorescence of a cell is dependent on the number of calcium ions bound to the indicator. if we denote [Ca^2+^] as the amount of free calcium in the cell and [B] as the number of proteins in the cell, we can use the binding/unbinding dynamics, coupled with the entry/exit dynamics of [Ca^2+^] in the cell to determine the fluorescence level at any given time. Specifically, we use the nonlinear diffusion of [Ca^2+^]

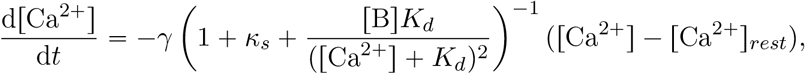

where [Ca^2+^]_*rest*_ represents the baseline free [Ca^2+^], *γ* is the [Ca^2+^] diffusion constant, *κ_s_* is the endogenous [Ca^2+^] binding ratio, and *K_d_* is the protein binding affinity constant [67, 68]. As *γ* is a function of the volume-to-surface area, we use a different *γ* value for dendrite dynamics as for dynamics in the soma [99]. While this model permits simulation of the [Ca^2+^]concentration over time, the model does not include the on/off time constants *τ_on_* and *τ_off_* that describe how long it takes for the bound proteins to become active. We can model this effect, as in [68], by convolving with a double-exponential function

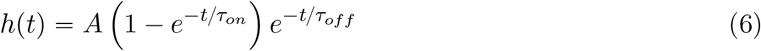

where the amplitude *A* and the time constants *τ_on_* and *τ_off_* can be fit to the particular protein kinetics. The final step in simulating the fluorescence time-traces is to convert the calcium concentrations to fluorescence levels. For this task, we use the Hill equation

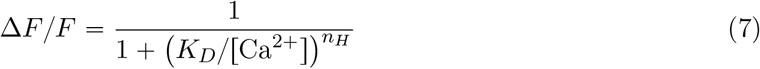

where the parameters *K_D_* and *n_H_* have been measured in the literature (specifically [70], Table 1), and the absolute florescence is

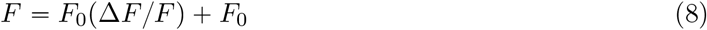

where the baseline fluorescence *F*_0_ can be tuned to the protein statistics.

### 4.4 Optics simulation

The optics module consists of modeling the shape and intensity of the point-spread function (PSF) within the scanned tissue. For computational purposes, we assume the shape of the PSF is constant across the scanned volume and only the amplitude is modulated. We estimate the PSF within the scanned tissue by propagating a specified field through the simulated tissue across the field of view.

We describe the scalar field at the front aperture of the objective lens as a Gaussian with a circular aperture and spherical phase:

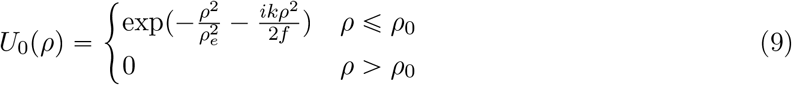

where *U*_0_ is the scalar field, 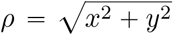 is the polar position, *k* is the wavenumber, *ρ*_0_ is the radius of the objective lens, *ρ_e_* is the radius of the excitation beam, and *f* is the focal length of the objective lens. The wavefront is multiplied by any additional specified aberrations due to the microscope or the sample:

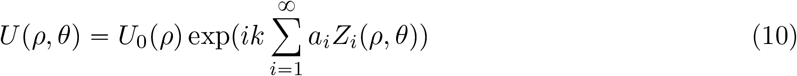

where *θ* is the polar angle, *a_i_* are the Zernike coefficients, and *Z_i_* are the Zernike polynomials. By default, only spherical aberration approximating the contribution of the refractive index mismatch of the sample and astigmatism approximating the contribution of offset scanning galvanometers are included.

The field *U*(*ρ, θ*) is propagated through the sample to the focal plane along a 2D grid of positions within a simulated refractive index volume *δn*. The volume *δn* is generated from the simulated vasculature and a 3D Gaussian process with a weight distribution approximating the refractive index distribution of mouse cortical tissue (see Supp Fig. 10) [100]:

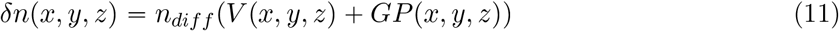

where *V* is the vasculature and *GP* is the smooth Gaussian Process representing the optical properties in the non-vasculature areas. The vasculature provides the bulk of the long range refractive index shifts in the simulation, while the Gaussian process approximates the local shifts.

The Fresnel diffraction integral is used to estimate the field throughout the volume, and the split-step beam propagation method [101] is used to apply the effects of inhomogeneity within the volume. The simulated phase-difference volume is summed into optical phase masks corresponding to each propagation step:

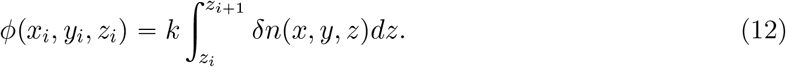

This quantity is multiplied after each optical propgation step as

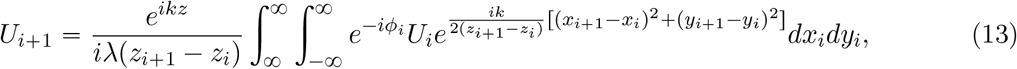

where *ϕ_i_* = *ϕ*(*x_i_,y_i_,z_i_*) is the optical phase mask and *U_i_* = *U*(*x_i_,y_i_,z_i_*) is the scalar field at each position. The resultant 3D field generated by the propagation is then used to calculate the two-photon PSF:

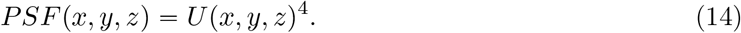

The aberrations caused by the phase differences approximate the effects of wavefront distortions caused by refractive index inhomogeneity within the imaged sample [102, 103]. The two-photon PSFs at each location across the field of view are averaged to obtain the PSF to be scanned through the simulation, and the summed intensity of the PSFs across the field are used to generate an intensity scaling mask for scanning. For runtime considerations, the PSF near the focal plane is sampled at the resolution of the volume while the out of focus PSF and scaling mask is sampled at a reduced resolution.

For alternative optical setups, we adjust the input field *U*_0_ accordingly. For a low numerical aperture excitation beam, *ρ_e_* is reduced, and for a Bessel beam excitation *U*_0_ is replaced with an excitation ring. See supplementary information for more details.

An additional optical mask is also calculated by estimating the reduction in signal from absorption of the collected light by the vasculature. The collected light at each scanned position is reduced by a collection cone corresponding to the simulated collection objective numerical aperture:

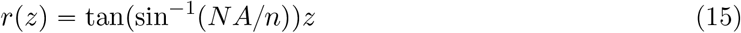

where *r*(*z*) is the collected cone radius as a function of depth, and:

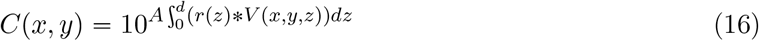

where *C* is fraction of light collected, *d* is the tissue depth, and *A* is the adjusted light absorbance of light emitted from GFP normalized by the arterial blood absorbance factor. This absorbance mask is multiplied to the optical excitation mask to give the combined spatial signal scaling mask.

### 4.5 Scanning *in silico*

The final module takes the generated volume, the generated PSF and time-traces, and generates the TPM output frames. The first step here is to use the time-traces and fluorescence distribution for each neuron to “color in” the corresponding volume with the current fluorescence level for that neuron. Similarly, the background level is set by repeating this process with the neuropil. The PSF is then convolved with the current volume, and the result is masked with the optical path mask to create an initial image.

To simulate motion in the movie, we select a portion of this initial frame to treat as the entire image. The starting position (upper left corner) for the with-motion frame is moved according to a small +/-0.5*μ*-m jitter with occasional larger jumps (up to 2-3*μ* m). Options to include per-line motion and shearing are also implemented by choosing different sub-sections of each row as the with-motion frame is extracted from the larger motionless frame. This frame represents the fluorescence level at each point in the sampled image. To obtain the actual electrical signals sampled by the TPM device, we apply a noise model that simulates the number of photons incident on the array (modeled as Poisson) followed by an electrical noise model that is Gaussian, with increasing mean and variance with larger numbers of incident photons. If λ is the true florescence for a pixel, *x* is the number of incident photons, and *y* is the measured electrical signal, the noise model can be expressed as

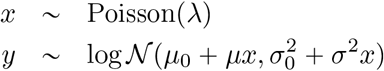

where *μ*_0_ and 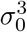 are the baseline noise mean and variance (with no photons), and *μ* and *σ*^2^ are the parameters controlling how the measurement mean and variance grow with increased incident photons.

As a final step, we simulate the analog-to-digital accumulators’ property where photons arriving in one pixel’s accumulation time can cause an analog shape that bleeds through to the accumulation for the next pixel (Fig. 12). We simulate this effect by noting that if a photon arrives early in the sample period, then the analog PMT response *g*(*t*) is completely inside of the sample period and no bleed-through occurs. On the other hand, if the photon arrives within Δ of the end of the sampling period, where Δ is the temporal extent of *g*(*t*) (Fig. 12), then the tail end of *g*(*t*) that continues beyond the end of the period is integrated into the next sample. The probability of a given bleed-through level for one photon can thus be quantified as

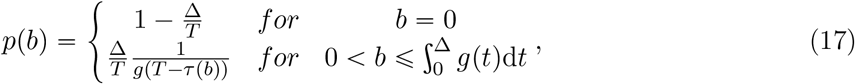

where *τ*(*b*) represents the delay *τ* that is needed to result in a given bleed-through *b*. Since the relationship between *b* and *τ*,

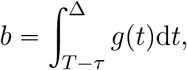

is monotonically increasing when *g*(*t*) ⩾ 0 for all *t*, *τ*(*b*) is a well defined function. Since photon arrivals are approximately independent, the bleed-through probability distribution for multiple photons is the convolution of the distribution for a single photon. The resulting statistical model then takes a random fraction (uniformly chosen between zero and 50%) of each pixel with probability 0.2, and adds that amount to the next pixel,

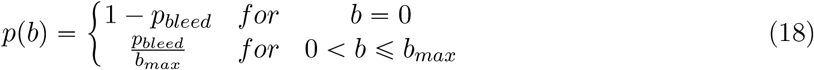

### Hemoglobin absorption

To calculate the absorption due to hemoglobin, we assume default concentrations pf 150mg/ml Hb, 64500 g/mol Hb, 2.9 (abs/*μ*m)/(mol/L) in units of abs/*μ*m. The absorbance is then calculated using Scott Prahl’s Hb curve [104] and eGFP emission spectrum [105].

### Estimation of per-trace noise variance

To estimate the noise variance for each time-trace, we begin with the basic per-pixel noise model

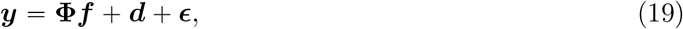

where the noise is heteroskedastic in that the variance is proportional to the mean

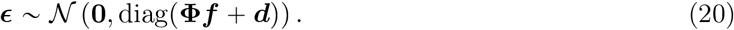

The least-squares estimate of the activations under the imperfect spatial profiles 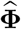 is

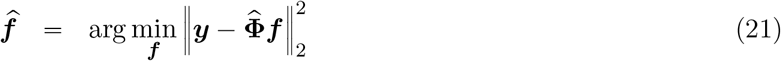

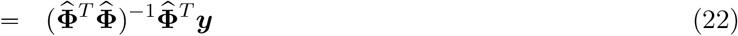

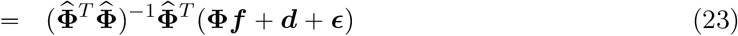

The covariance of this estimate is then

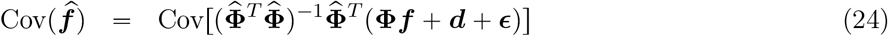

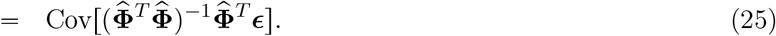

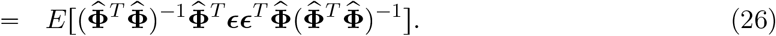

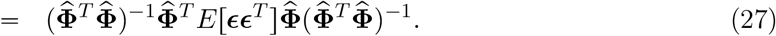

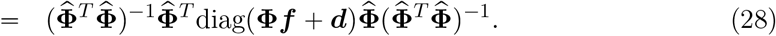

### Local correlation calculation

To calculate the local correlations, V1 two-photon recordings and simulations were motion corrected using correlation-based rigid motion correction [106]. A 1500-frame subsection of each dataset over a 250 × 250 pixel area was extracted. For each pixel, the Pearson correlation between it’s fluorescence activity and that of each of the neighboring pixels in a 51 × 51 pixel square neighborhood (up to 25 pixels away in each direction) were calculated. The results were averaged over all pixels (to create the mean images) and histograms were created to depict the spread for correlations along the fast-scan direction.

### Calculation of auxiliary time-traces

To calculate the auxiliary, noisy “ground truth” time traces, we consider the movie frames ***y***_*t*_ for *t* = 1…*T* and the calculated ground-truth spatial profiles ***X*** = [*x*_1_,…, *x_N_*]. The noisy time trace estimates are then calculated via the least-squares estimation procedure at each time-step *t*

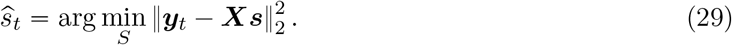

## Acknowledgements

The authors would like to thank Thomas Macrina and Sebastian Seung for helpful discussions of anatomy and electron microscopy data, and Chris Schaffer and Mohammad Haft-Javaherian for the use of and helpful discussions pertaining to their vasculature data. Moreover, the authors would like to thank Ben Scott, Sam Wang, Sue Ann Koay, Brian DePasquale and Dylan Rich For their insightful thoughts and comments throughout this project. A.S.C. and J.W.P. were supported by Simons Collaboration on the Global Brain (SCGB AWD543027), and a U19 NIH-NINDS BRAIN Initiative Award (5U19NS104648). A.S. was supported by NIH grant 1U19NS104648. D.W.T. was supported by NIH grants R01MH083868 and 1U19NS104648, and the Simons Collaboration on the Global Brain (SCGB 543051).

## Author Contributions

A.S.C. and A.S. designed and coded the bulk of the simulation software and performed the ensuing analysis. J.L.G. helped design the scanning noise model. A.S.C., A.S., J.W.P. and D.W.T. wrote the manuscript. J.W.P. and D.W.T. supervised the project.

## Competing Financial Interests

The authors declare no competing financial interests.

## 4.6 Supplementary tables

**Supplementary Table 1:** Anatomical parameters used for *in-silico* simulation of neural activity in layer II/III of mouse primary visual area V1. Values for each parameter were either directly found in the literature or estimated from published data (entries with a †). The third column indicates whether these parameters were set directly in NAOMi, or were fit indirectly by setting other simulation parameters. In the latter cases, the measured values from a simulated NAOMi volume are shown for comparison, indicating that the simulated anatomy matches measured anatomical statistics.

| Anatomical: |  |  |  |  |  |
| --- | --- | --- | --- | --- | --- |
| Parameter | Lit. Val. | Fit type | NAOMi Val. | Unit | Refs |
| Neural density | 9.20E+04 | Direct | – | mm<sup>-3</sup> | [58] |
| Axonal diameter | 0.3 | Direct | – | μm | [58] |
| Fraction vasculature | 0.01-0.04 | Indirect | 0.032 | – | [58, 74, 75] |
| Fraction cell bodies | 0.12 | Indirect | 0.135 | – | [58] |
| Fraction neuropil | 0.84 | Indirect | 0.833 | – | [58] |
| Fraction dendrites* | 0.294 | Indirect | 0.223 | – | [58] |
| Fraction other (fluorescing)* | 0.401 | Indirect | 0.33 | – | [58] |
| Fraction other (not fluorescing)* | 0.293 | Indirect | 0.28 | – | [58, 76] |
| Vessel radius (capillary) | 2.00E+00 | Direct | – | μm | [56] |
| Vessel radius (penetrating) | 10 (9,11) | Direct | – | μm | [56] |
| Vessel radius (surface) | 4-30 | Direct | – | μm | [56] |
| Vessel length | 50 | Indirect | 50 | μm | [56] |
| Vessel orientation | Uniform | Indirect | Uniform | – | [56] |
| Vascular density | 1-3 | Indirect | 2 | % | [56, 58, 74, 75] |
| Penetrating vessel density | 30<sup>†</sup> | Direct | – | mm<sup>-2</sup> | [56] |
| Somatic volume* | 1.80E+03<sup>†</sup> | Indirect | 1.80E+03 | μm<sup>3</sup> | [77] |
| Nuclear volume* | 800<sup>†</sup> | Indirect | 800 | μm<sup>3</sup> | [77] |
| Cytoplasm volume* | 1000<sup>†</sup> | Indirect | 1000 | μm<sup>3</sup> | [77] |
| Somatic radius* | 6.25-7.5<sup>†</sup> | Indirect | 7.5 | μm | [60, 77] |
| Nuclear radius* | 6<sup>†</sup> | Indirect | 6 | μm | [77] |
| Fraction cell bodies | 0.12 | Indirect | 0.135 | – | [58] |
| Basal dendrite endings | 23-25<sup>†</sup> | Direct | – | – | [59–61] |
| Basal dendrite diameter | 0.7 | Direct | – | μm | [78] |
| Basal dendrite length | 100-160<sup>†</sup> | Indirect | 105 | μm | [59–61] |
| Apical dendrite endings | 6-20<sup>†</sup> | Direct | – | – | [59, 60] |
| Apical dendrite diameter | 1-2 | Direct | – | μm | [79] |
| Rall exponent | 1.5 | Direct | – | – | [107] |
| *Adjusted for shrinkage: 31% [86],<sup>†</sup>Value estimated from data in the literature |  |  |  |  |  |

**Supplementary Table 2:** Fluorescence parameters used for *in-silico* simulation of neural activity in layer II/III of mouse primary visual area V1. Values for each parameter were either directly found in the literature or estimated from published data (entries with a †). The third column indicates whether these parameters were set directly, or were fit indirectly by setting other simulation parameters. In the latter cases, the measured values simulated fluorescence traces are shown for comparison, indicating a good match between the NAOMi simulation and known activity statistics.

| Fluorescence model: |  |  |  |  |  |
| --- | --- | --- | --- | --- | --- |
| Parameter | Anat. Val. | Fit type | NAOMi Val. | Unit | Refs |
| GCaMP6f binding affinity $K_d$ | 290 | Direct | – | nMol | [70] |
| Baseline $\text{Ca}^{2+}$ concentration | 50.00 | Direct | – | nMol | [68] |
| $\text{Ca}^{2+}$ axon diffusion constant $\gamma$ | 2800 | Direct | – | $\text{s}^{-1}$ | [67, 80, 83] |
| Endogenous $\text{Ca}^{2+}$ binding ratio $k_s$ | 100,110 | Direct | 110 | AU | [67, 80–82] |
| $\text{Ca}^{2+}$ diffusion constant $\gamma$ | 1800 | Data fit | 292.3 | $\text{s}^{-1}$ | [67, 80, 81] |
| Onset time constant $\tau_{on}$ | – | Data fit | 98.62 | AU | – |
| Decay time constant $\tau_{off}$ | – | Data fit | 0.85 | AU | – |
| GCaMP6f Hill eqn. exponent $n_h$ | 2.7 | Direct | – | AU | [70] |
| GCaMP6f Hill eqn. amplitude | 25.2 | Direct | – | F | [70] |
| Indicator concentration | 10-200 | Direct | 10 | $\mu\text{M}$ | [68, 84, 85] |

## 4.7 Supplementary figures

**Supplementary Figure 1:**
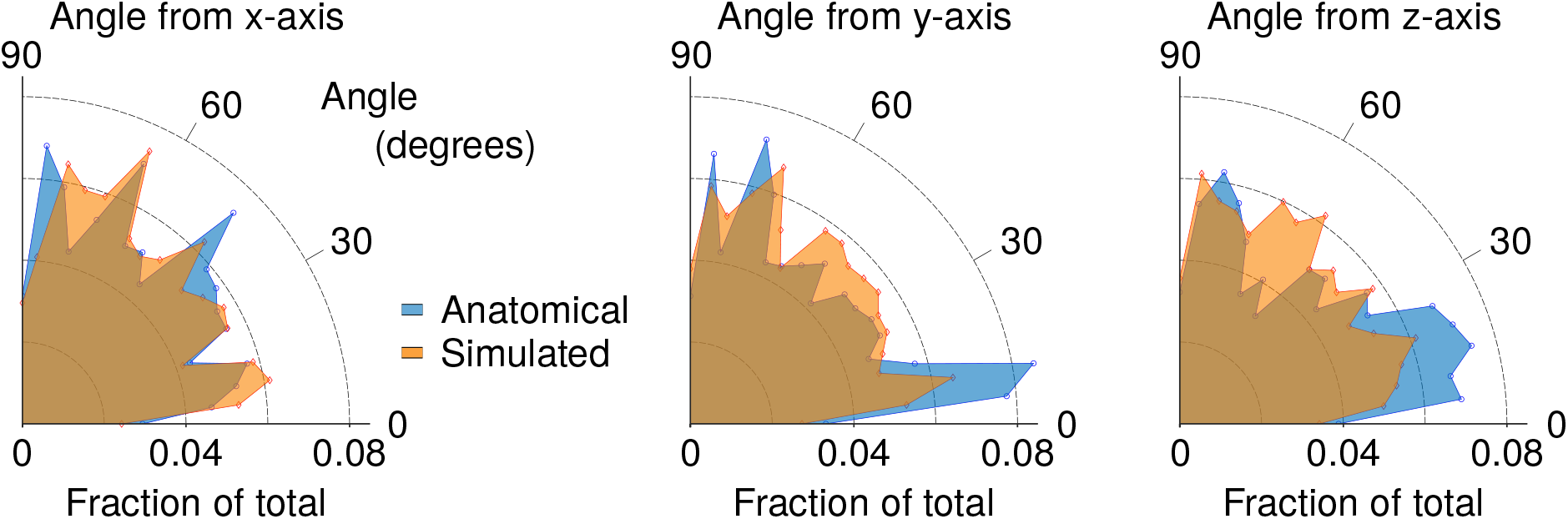
Histograms comparing the distribution of vasculature orientation (angles from the x-y- and z-axes).

**Supplementary Table 3:** Optical parameters used for *in-silico* simulation of two-photon microscopy scanning.

| Optical: |  |  |  |
| --- | --- | --- | --- |
| Parameter | Value | Unit | Refs |
| Refractive index (tissue average) | 1.35 | – | [108] |
| Refractive index (SD) | 0.02 | – | [77, 108] |
| Scatter size | [0.51 1.56 4.52 14.78] | $\mu\text{m}$ | [77, 100], See Methods |
| Scatter weight | [0.57 0.29 0.19 0.15] | – | [77, 100], See Methods |
| Wavelength $\lambda$ | 920 | nm | [109] |
| System aberrations (astigmatism) | 0.1 | $\lambda$ | See Methods |
| System aberrations (spherical) | 0.12 | $\lambda$ | See Methods |
| Hemoglobin absorption | $0.00674 \cdot \log(10)$ | – | [104], See Methods |
| Objective NA | 0.8 | – | [110] |
| PMT QE | 0.4 | – | [111] |
| eGFP Quantum Yield | 0.6 | – | [109] |
| $\Delta$, baseline $F_0$ | 2 | GM | [109] |
| $\Delta$, saturation $F_{max}$ | 35 | GM | [109] |
| Sech pulse shape | 0.588 | – | [112] |
| Ti:S rep rate | 80 | MHz | [112] |
| Ti:S pulse width | 140 | fs | [112] |

**Supplementary Table 4:**
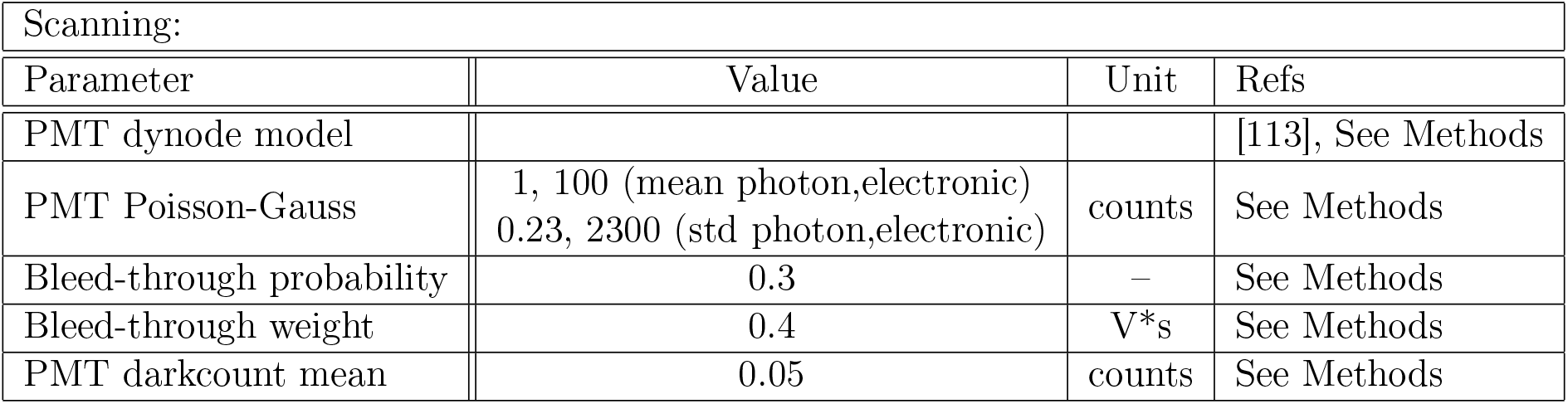
Scanning parameters used for *in-silico* simulation of two-photon microscopy scanning.

| Scanning: |  |  |  |
| --- | --- | --- | --- |
| Parameter | Value | Unit | Refs |
| PMT dynode model |  |  | [113], See Methods |
| PMT Poisson-Gauss | 1, 100 (mean photon,electronic)<br>0.23, 2300 (std photon,electronic) | counts | See Methods |
| Bleed-through probability | 0.3 | – | See Methods |
| Bleed-through weight | 0.4 | V*s | See Methods |
| PMT darkcount mean | 0.05 | counts | See Methods |

**Supplementary Figure 2:**
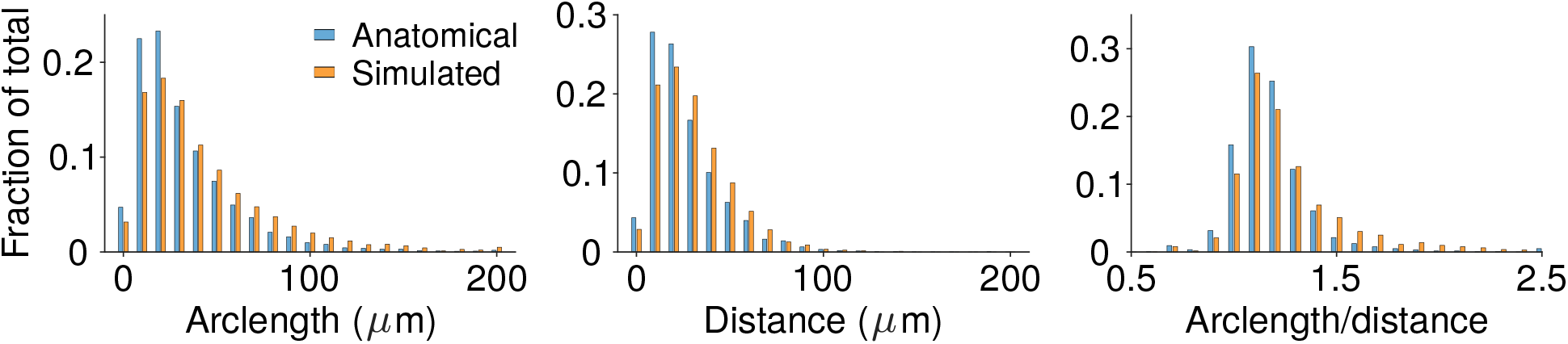
Histograms comparing the lengths of vasculature.

**Supplementary Table 5:** Detailed fractional volume of components

| Other parameters: |  |  |  |  |  |
| --- | --- | --- | --- | --- | --- |
| Parameter | Anat. Val. | Fit type | NAOMi Val. | Unit | Refs |
| Axonal diameter | 0.3 | Direct | – | um | [58] |
| Fraction vasculature | 0.01-0.04 | Indirect | 0.032 | – | [58, 74, 75] |
| Fraction cell bodies | 0.12 | Indirect | 0.135 | – | [58] |
| Fraction neuropil | 0.84 | Indirect | 0.833 | – | [58] |
| Fraction spines* | 0.118 | Indirect | 0 | – | [58] |
| Fraction dendrites* | 0.294 | Indirect | 0.223 | – | [58] |
| Fraction Axons* | 0.283 | Indirect | 0.33 | – | [58] |
| Fraction Glia* | 0.0924 | Indirect | 0.08 | – | [58] |
| Fraction extracellular | 0.2 | Direct | 0.2 | – | [76] |
| *scaled to full volume [86] |  |  |  |  |  |

**Supplementary Table 6:**
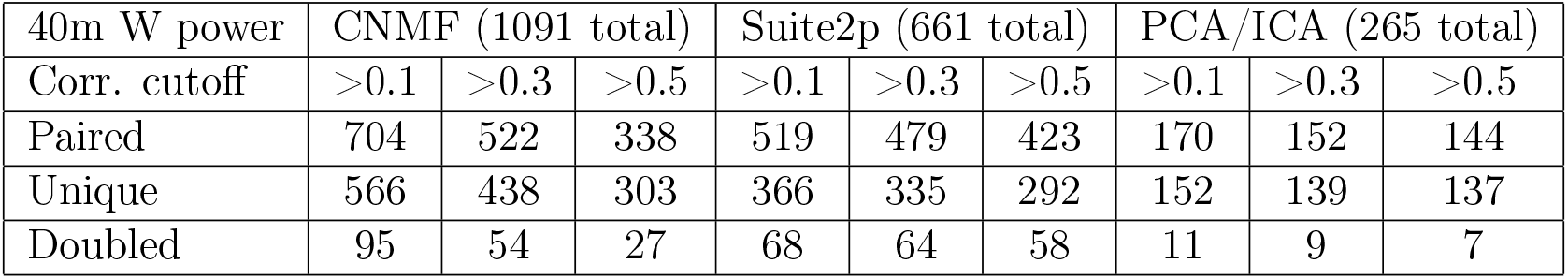
Results of automated calcium imaging video segmentation as applied to simulated data generated from NAOMi. Of the total number of components isolated, only a fraction (≈ 30% – 50%) unique, true cells in the scanned volume strongly matched the found components.

**Supplementary Table 7:** Results of automated calcium imaging video segmentation as applied to simulated data generated from NAOMi. While sets of found components largely overlapped between algorithms, each method’s design allowed for the extraction of slightly different cell activities.

| 40m W power | Unique Cells found |  |  |
| --- | --- | --- | --- |
| Corr. cutoff | >0.1 | >0.3 | >0.5 |
| CNMF (1091 total) | 566 | 438 | 303 |
| Suite2p (1091 total) | 366 | 335 | 292 |
| PCA/ICA (265 total) | 152 | 139 | 137 |
| CNMF Only | 238 | 69 | 28 |
| Suite2p Only | 43 | 26 | 17 |
| PCA/ICA Only | 13 | 2 | 3 |
| CNMF & Suite2p | 321 | 307 | 273 |
| CNMF & PCA/ICA | 144 | 136 | 134 |
| Suite2p & PCA/ICA | 134 | 137 | 134 |
| All algorithms | 137 | 135 | 132 |

**Supplementary Table 8:**
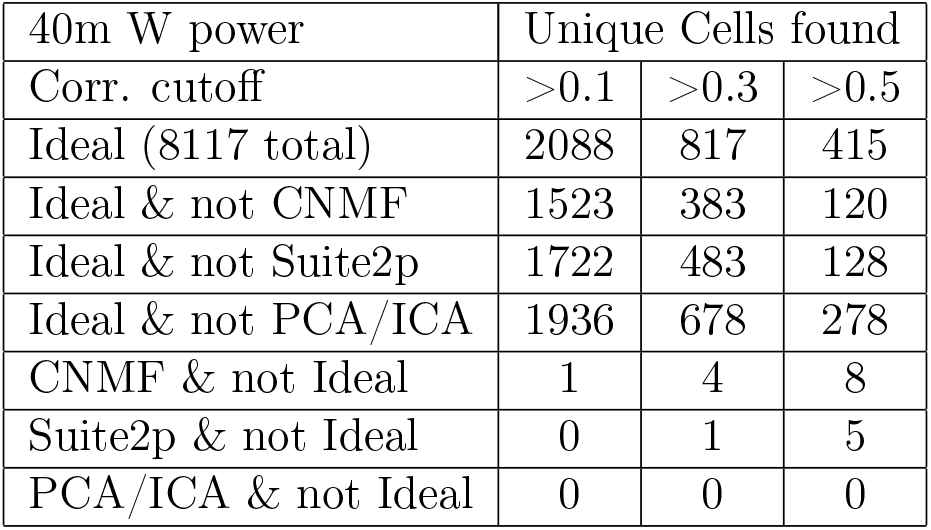
Results of automated calcium imaging video segmentation as applied to simulated data generated from NAOMi. De-mixing the data using oracle spatial profile knowledge allowed for isolating many more components, indicating that automated methods have room for improvement. Interestingly some algorithms, due to built in denoising not included in the ideal de-mixing, were able to isolate some cell time-traces more accurately.

| 40m W power | Unique Cells found |  |  |
| --- | --- | --- | --- |
| Corr. cutoff | >0.1 | >0.3 | >0.5 |
| Ideal (8117 total) | 2088 | 817 | 415 |
| Ideal & not CNMF | 1523 | 383 | 120 |
| Ideal & not Suite2p | 1722 | 483 | 128 |
| Ideal & not PCA/ICA | 1936 | 678 | 278 |
| CNMF & not Ideal | 1 | 4 | 8 |
| Suite2p & not Ideal | 0 | 1 | 5 |
| PCA/ICA & not Ideal | 0 | 0 | 0 |

**Supplementary Table 9:**
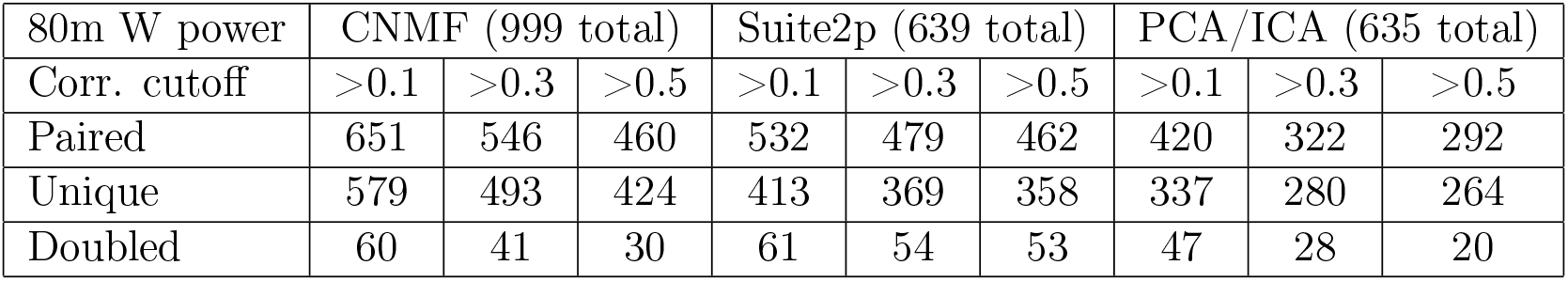
Results of automated calcium imaging video segmentation as applied to simulated data generated from NAOMi with 80m W laser power.

| 80m W power | CNMF (999 total) |  |  | Suite2p (639 total) |  |  | PCA/ICA (635 total) |  |  |
| --- | --- | --- | --- | --- | --- | --- | --- | --- | --- |
| Corr. cutoff | >0.1 | >0.3 | >0.5 | >0.1 | >0.3 | >0.5 | >0.1 | >0.3 | >0.5 |
| Paired | 651 | 546 | 460 | 532 | 479 | 462 | 420 | 322 | 292 |
| Unique | 579 | 493 | 424 | 413 | 369 | 358 | 337 | 280 | 264 |
| Doubled | 60 | 41 | 30 | 61 | 54 | 53 | 47 | 28 | 20 |

**Supplementary Table 10:** Results of automated calcium imaging video segmentation as applied to simulated data generated from NAOMi with 80m W laser power.

| 80m W power | Unique Cells found |  |  |
| --- | --- | --- | --- |
| Corr. cutoff | >0.1 | >0.3 | >0.5 |
| CNMF (999 total) | 579 | 493 | 424 |
| Suite2p (639 total) | 413 | 369 | 358 |
| PCA/ICA (635 total) | 337 | 280 | 264 |
| CNMF Only | 198 | 118 | 85 |
| Suite2p Only | 46 | 24 | 22 |
| PCA/ICA Only | 90 | 38 | 24 |
| CNMF & Suite2p | 357 | 334 | 325 |
| CNMF & PCA/ICA | 261 | 247 | 243 |
| Suite2p & PCA/ICA | 240 | 242 | 240 |
| All algorithms | 237 | 231 | 229 |

**Supplementary Table 11:**
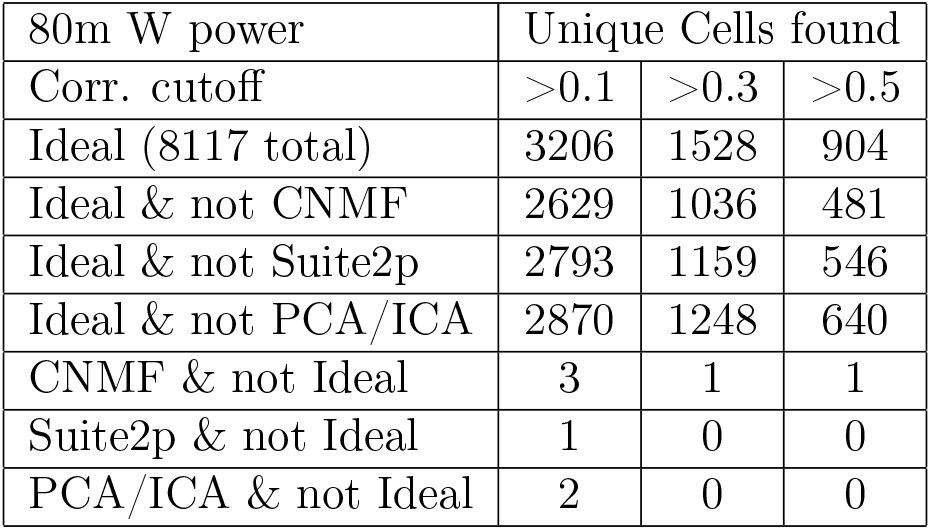
Results of automated calcium imaging video segmentation as applied to simulated data generated from NAOMi with 80m W laser power.

**Supplementary Figure 3:**
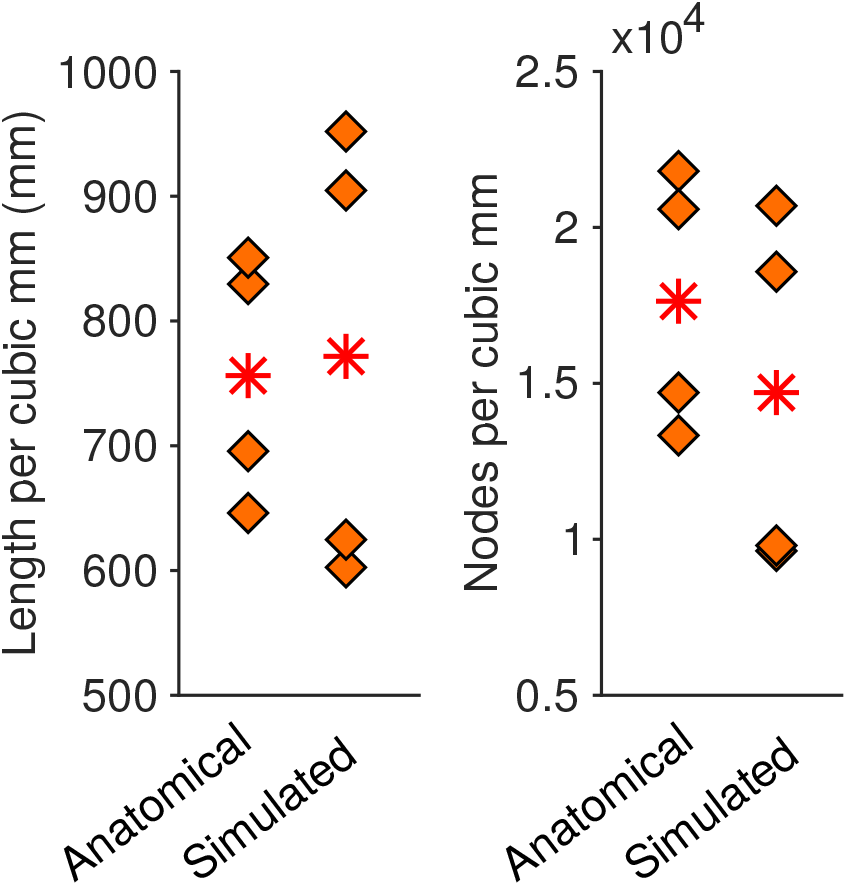
Comparison of overall length of the vasculature and the number of vasculature nodes within a cubic mm volume.

**Supplementary Figure 4:**
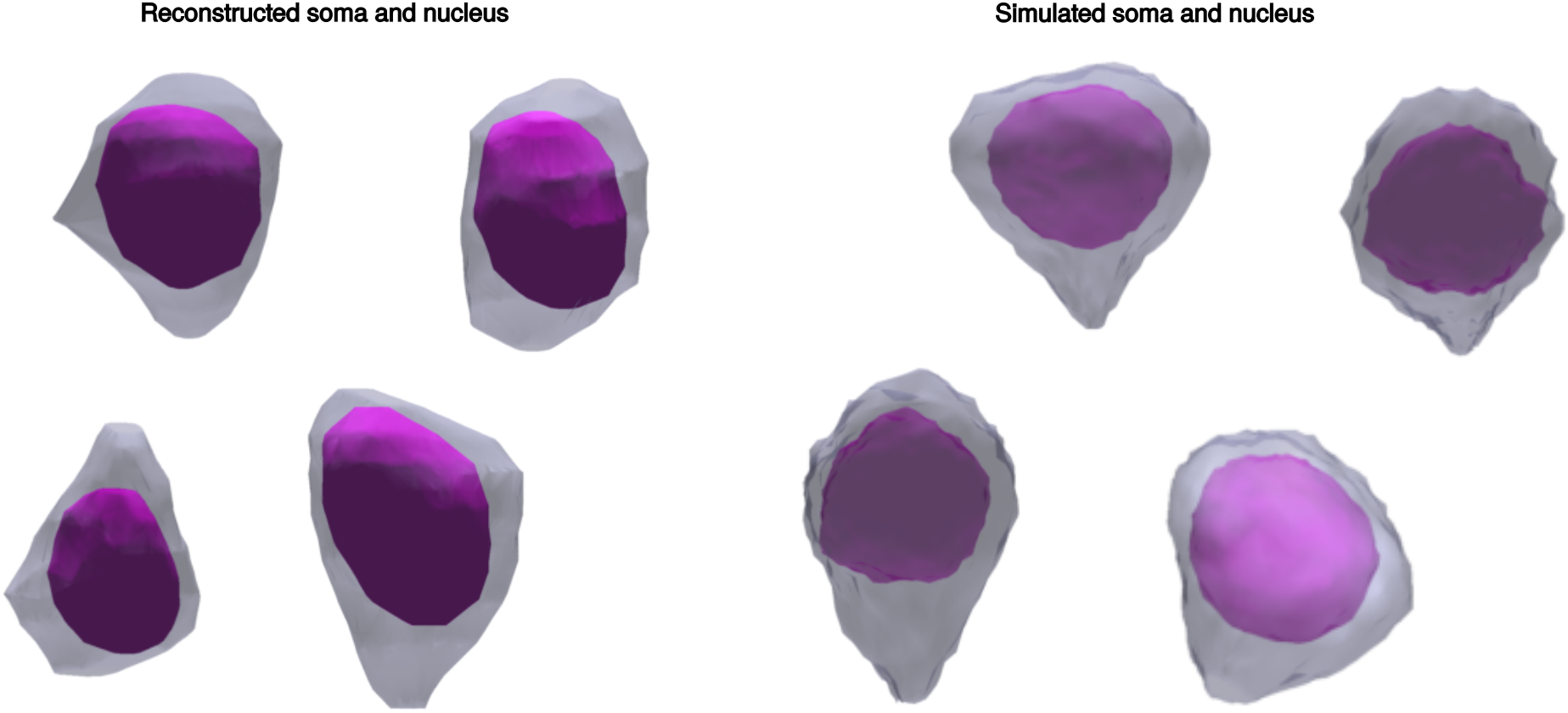
Examples of measured and simulated somas. Both measured somas (left) and simulated somas (right) have a bumpy cell wall with one end exhibiting the cone-shaped tightening characteristic of pyramidal cells. The nuclei of both sets of somas (blue shape inside of the red shape) are shrunken and smoothed versions of the exterior cell wall.

**Supplementary Figure 5:**
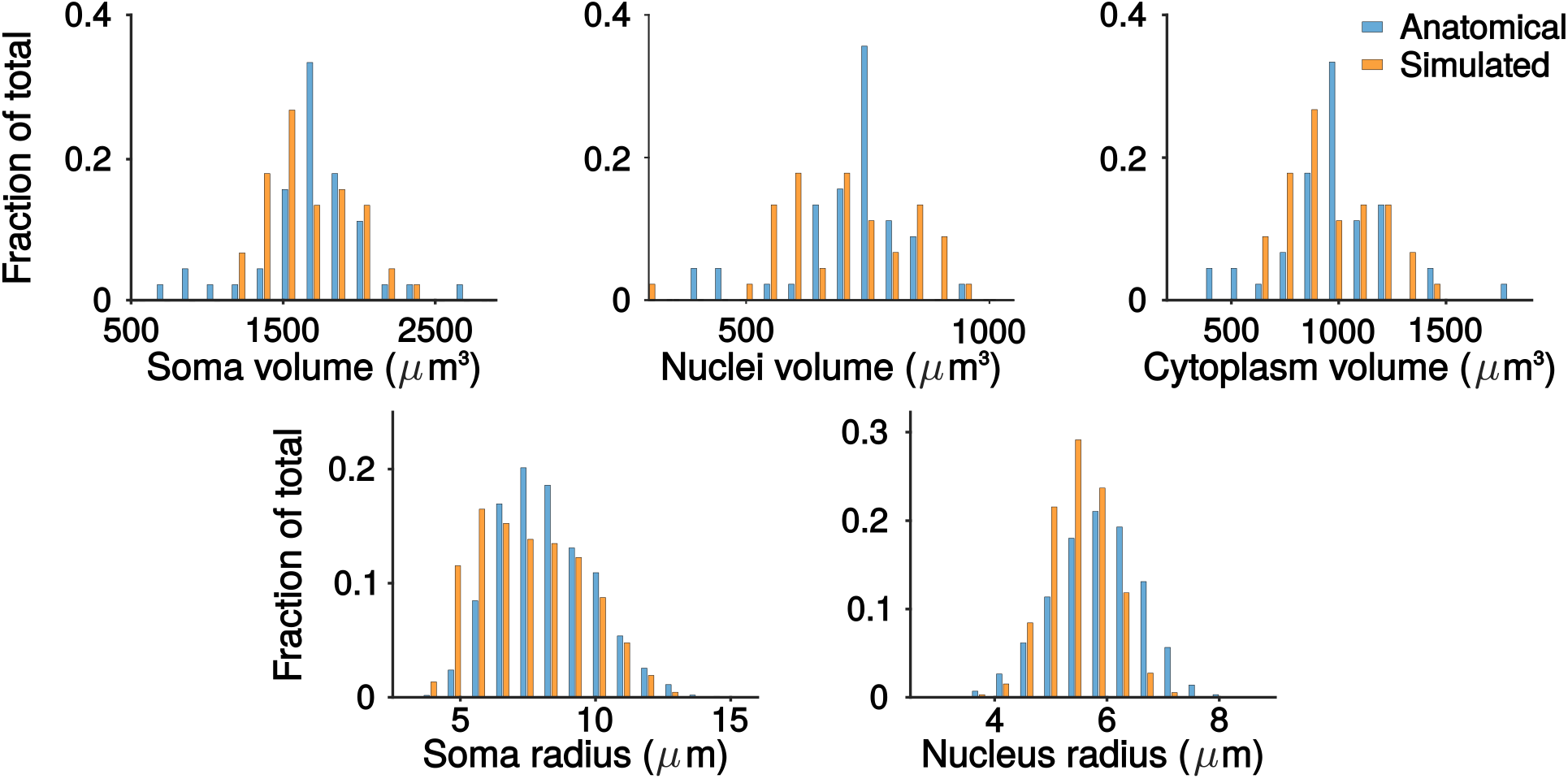
Histogram comparing simulated somas with measured anatomy.

**Supplementary Figure 6:**
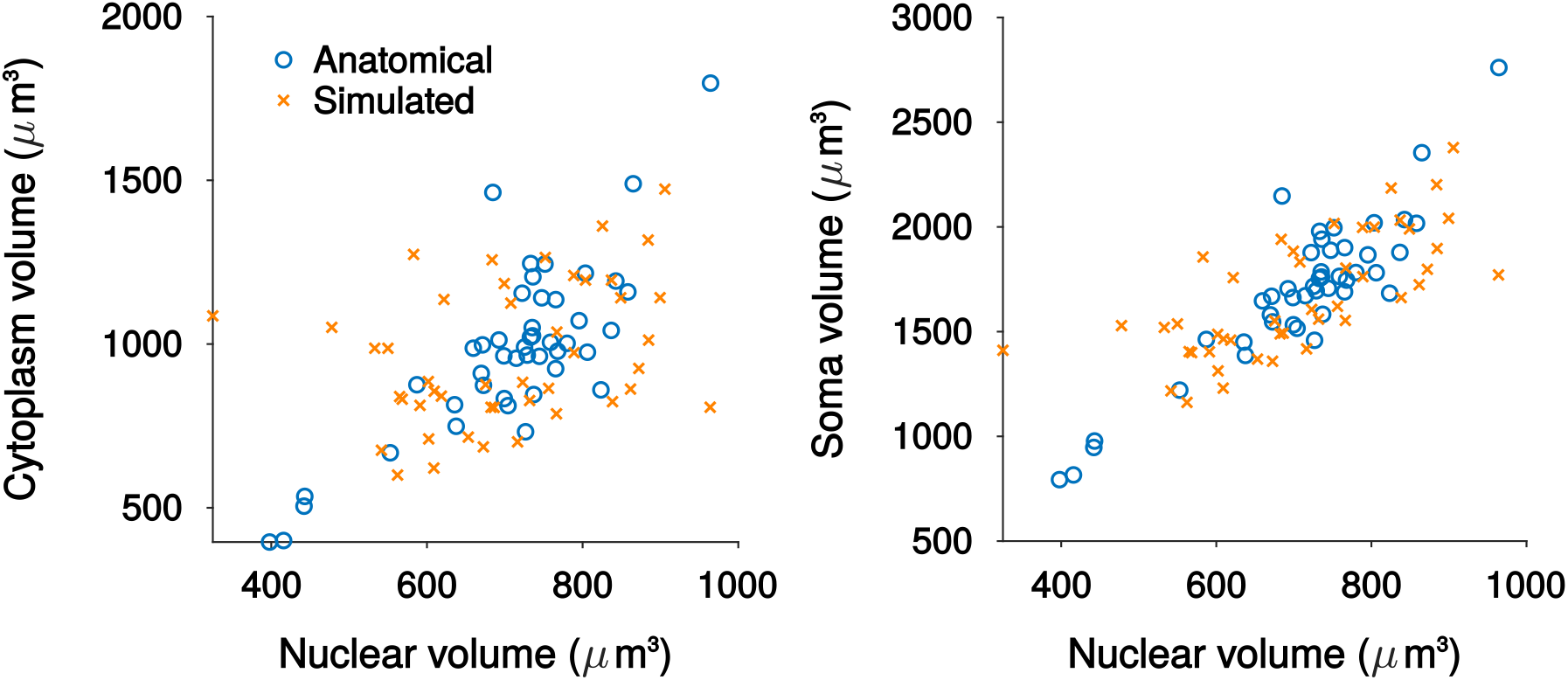
Scatter plots comparing the nucleus, soma and cytoplasm volumes for simulated neurons with measured anatomy from EM reconstructions.

**Supplementary Figure 7:**
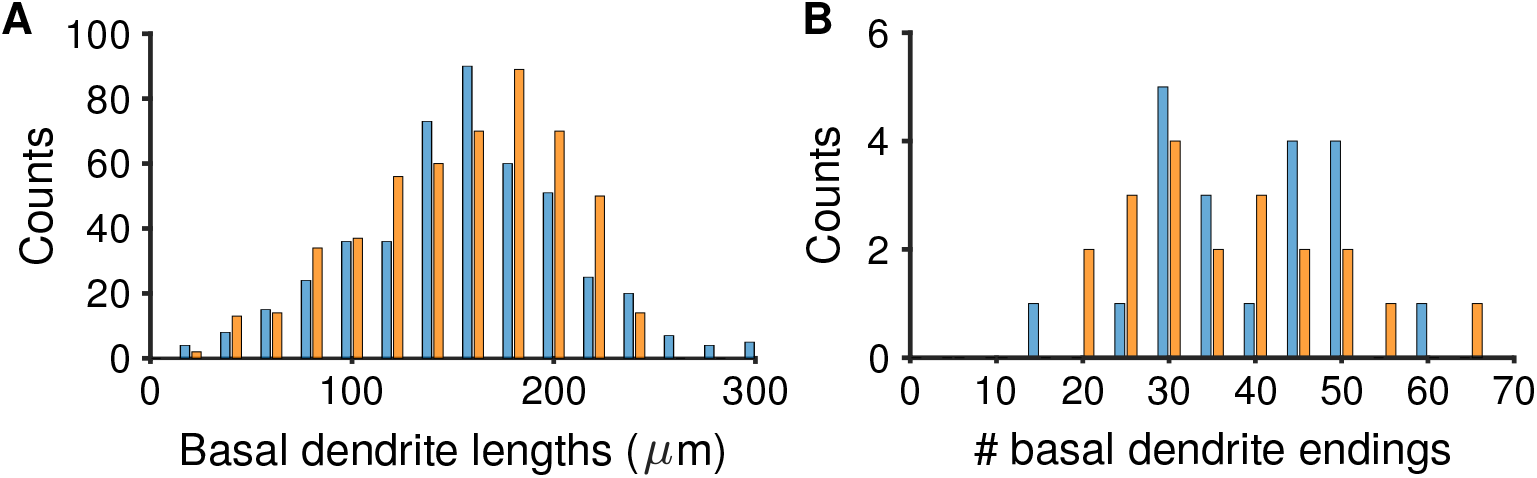
A: Histogram of total basal dendrite length per cell from measured [60] and simulated data. B: Histogram of total number of basal dendrite endings

**Supplementary Figure 8:**
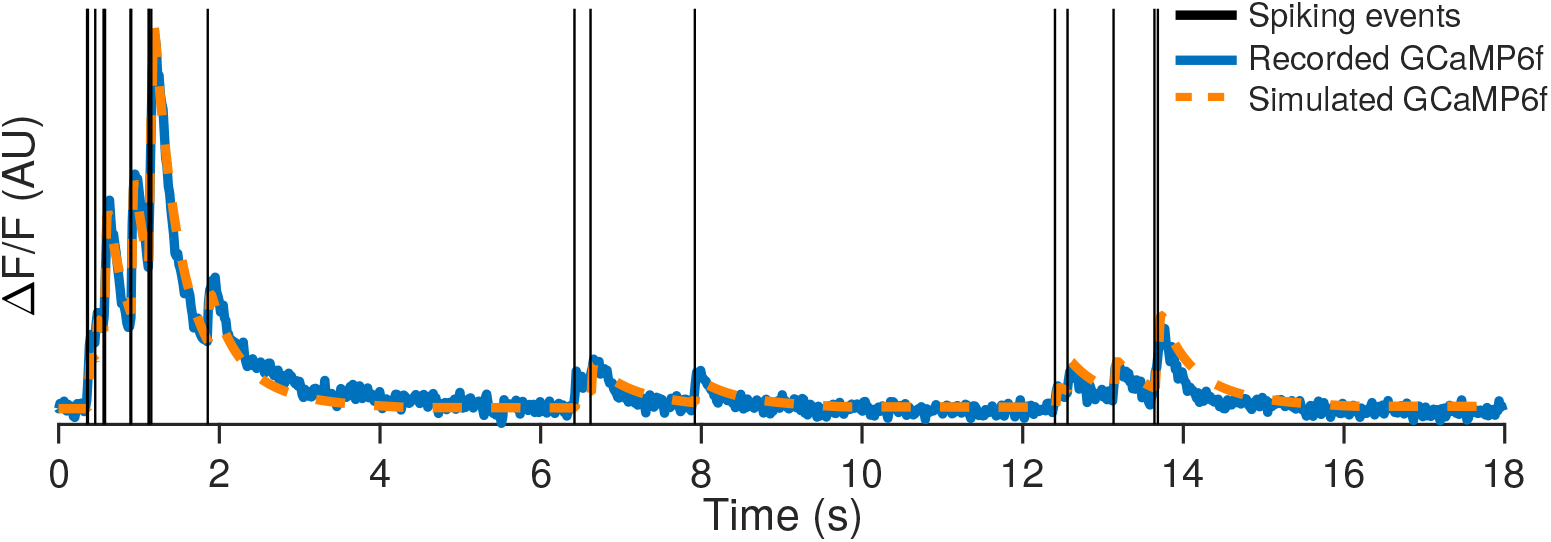
Simultaneously recorded spikes and two-photon fluorescence timecourse [114] along with the estimated fluorescence timecourse using a forward model of calcium response.

**Supplementary Figure 9:**
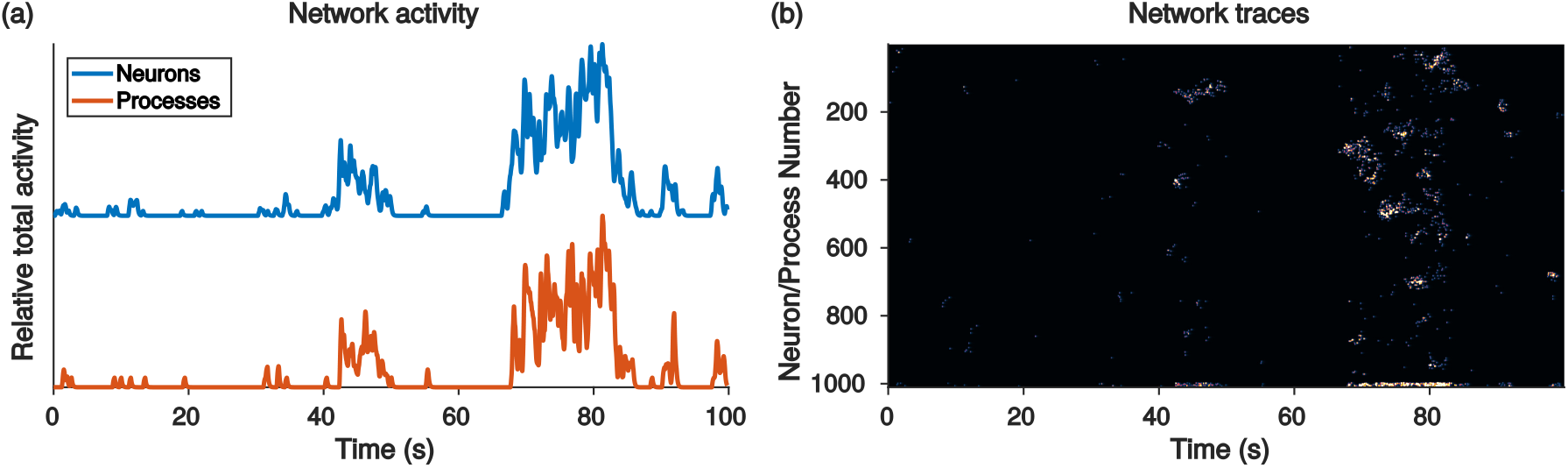
Example of a Hawkes point-process. (a) The total activity of the network is correlated with the total activity in the background processes. (b) The Hawkes process gives the network- and single neuron-bursting statistics common in many neural activity recordings.

**Supplementary Figure 10:**
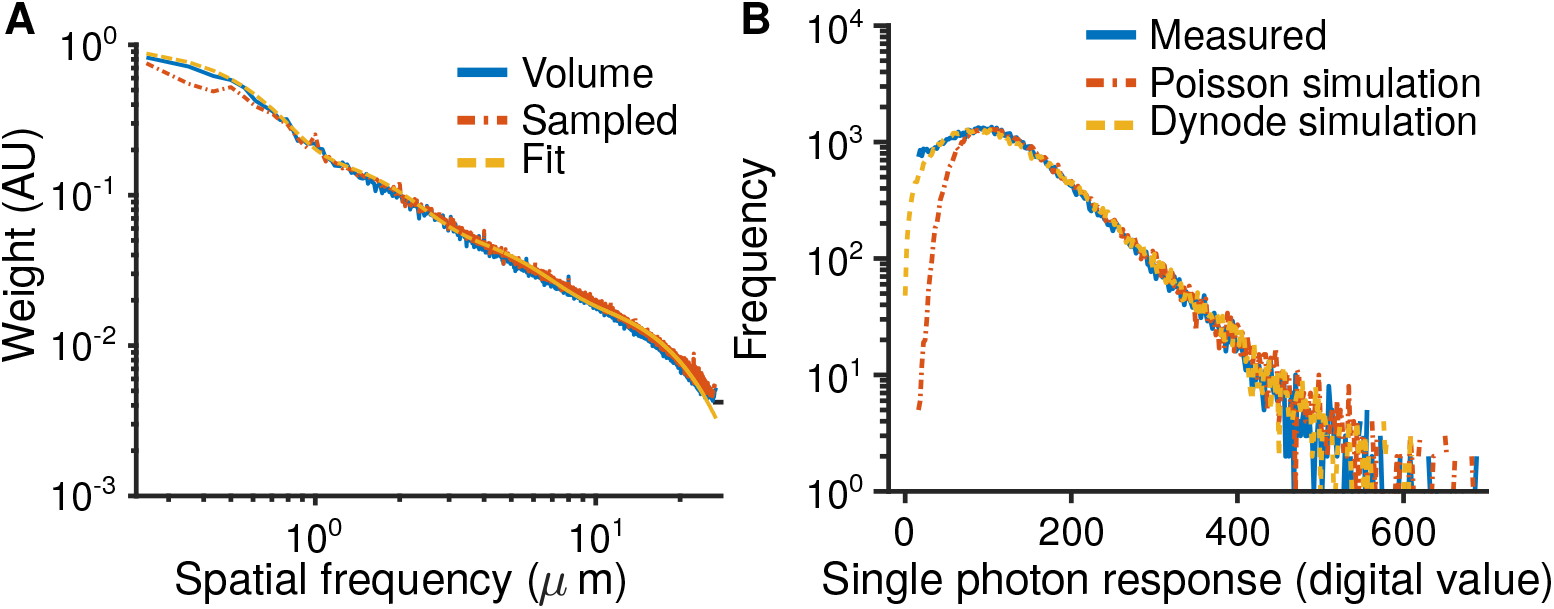
A: Spatial frequency weights of an EM volume of mouse visual cortex, the fit to a Gaussian mixture model (GMM), and weights from a 3D Gaussian process sampling using the GMM. B: Measured PMT single photon response digital count distribution as compared to a Poisson amplification model or a dynode amplification model.

**Supplementary Figure 11:**
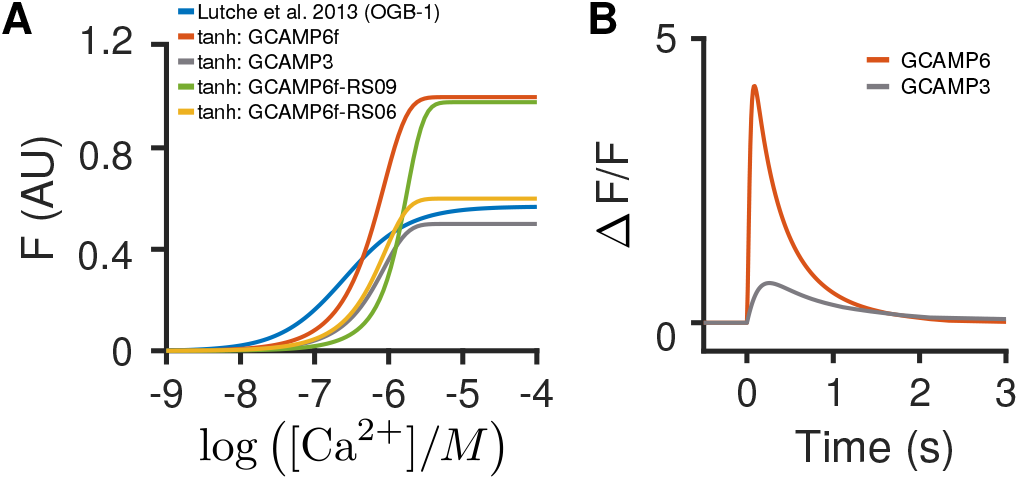
A: Example hill plots that can be used in the fluorescence simulation (relating calcium concentration to ΔF/F). The height and slope dictate the saturation and decay behavior. These curves were to the indicators in [71], using the parameters supplied therein. B: Example ΔF/F behavior for GcAMP6f and GcAMP6s in response to a ten-spike burst.

**Supplementary Figure 12:**
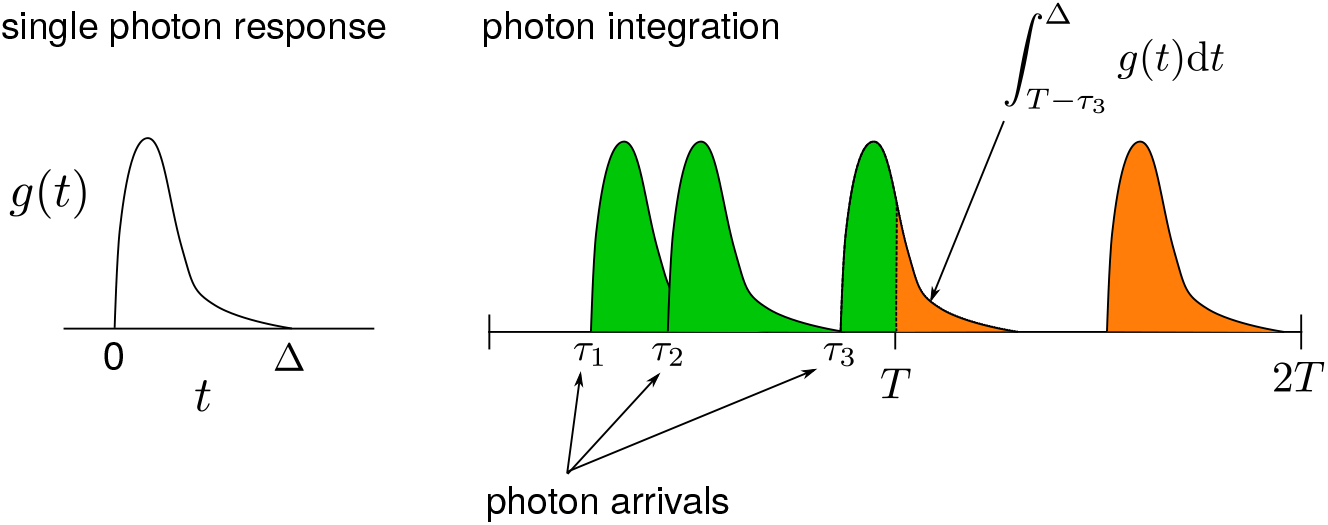
Bleed-through of photon responses during TPM electronic analog-to-digital conversion. Left: A single photon causes a response in the electronics that persists over a time frame Δ. Right: any photons that arrive within Δ of the end of the sampling period cutoff (every *T* seconds) have responses that are partially integrated into the current sample (green) and partially integrated into the next sample (orange).

**Supplementary Figure 13:**
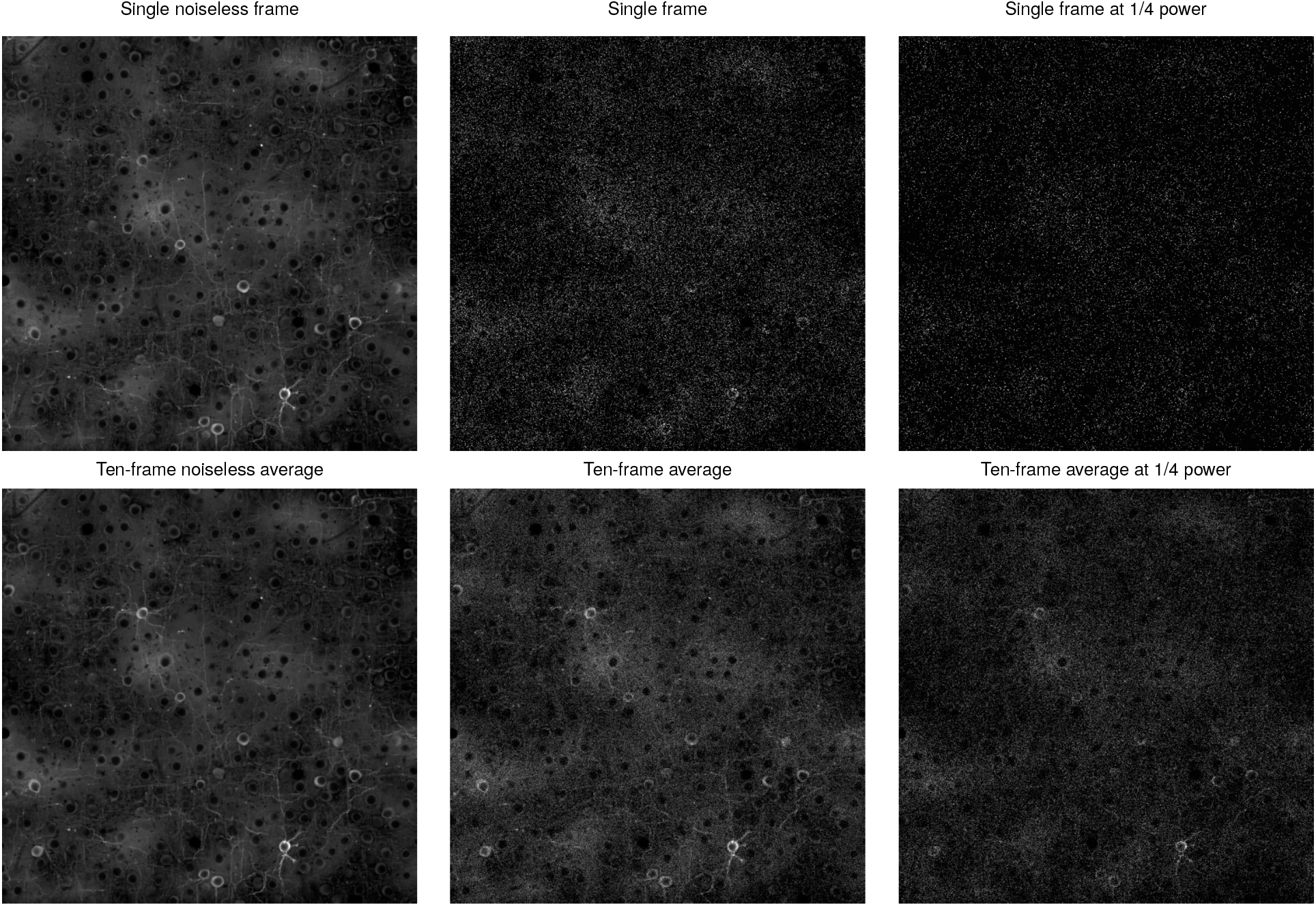
Example frames showing the effect of reducing the power of the scanning PSF.

**Supplementary Figure 14:**
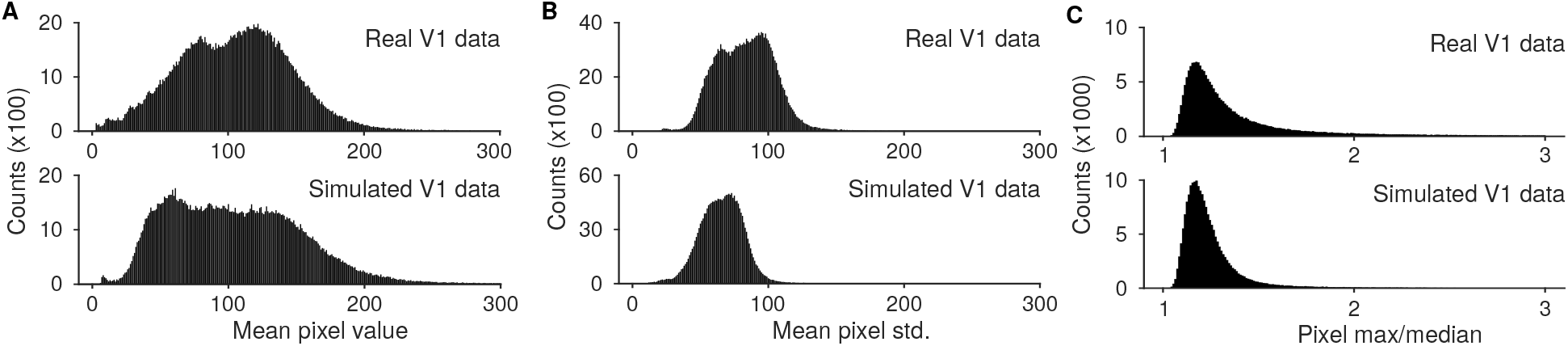
A: Histogram of values in the mean image over 20000 frames for both real and simulated V1 data. B: Histogram of standard deviations across the FOV over 2000 frames for both real and simulated V1 data. C: Histogram of the ratio of the maximum value to the median value (approximate estimte of activity) across all pixels in the FOV, calculated over 20000 frames.

**Supplementary Figure 15:**
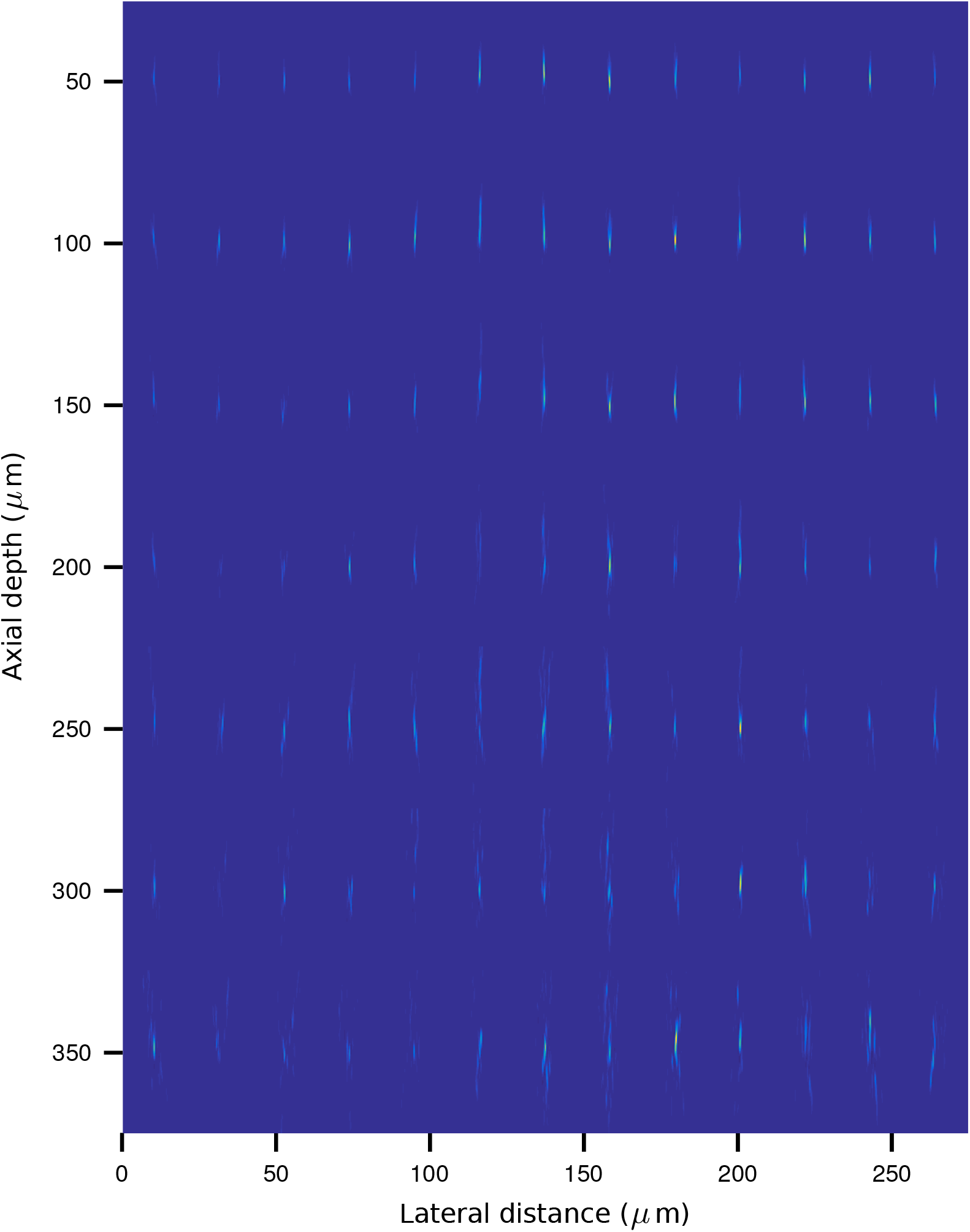
Simulated propagation of a point-spread function through layers of diffusing tissue using NAOMi. Simulated PSFs equally spaced across two ≈2000*μ*m cross sections of tissue, sampled at

**Supplementary Figure 16:**
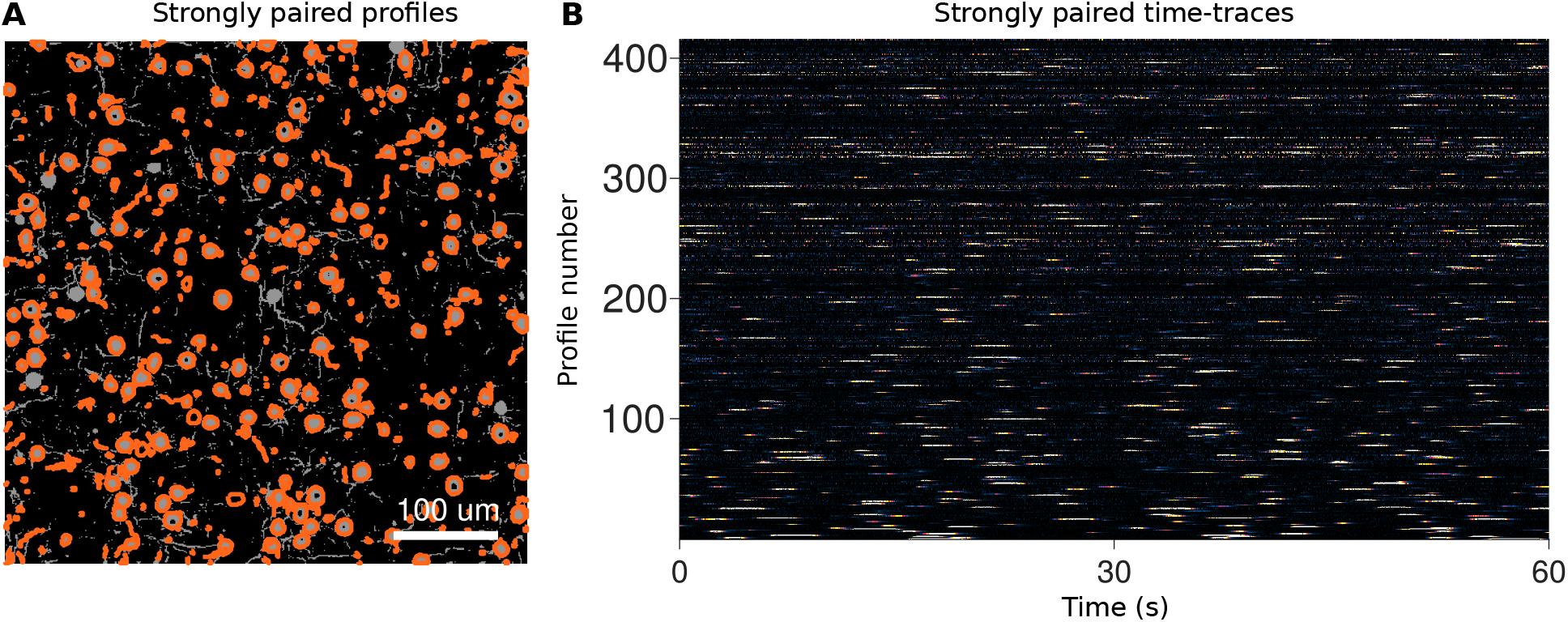
Results of using the ideal spatial components to de-mix the simulated TPM video with frame-by-frame least-squares.

**Supplementary Figure 17:**
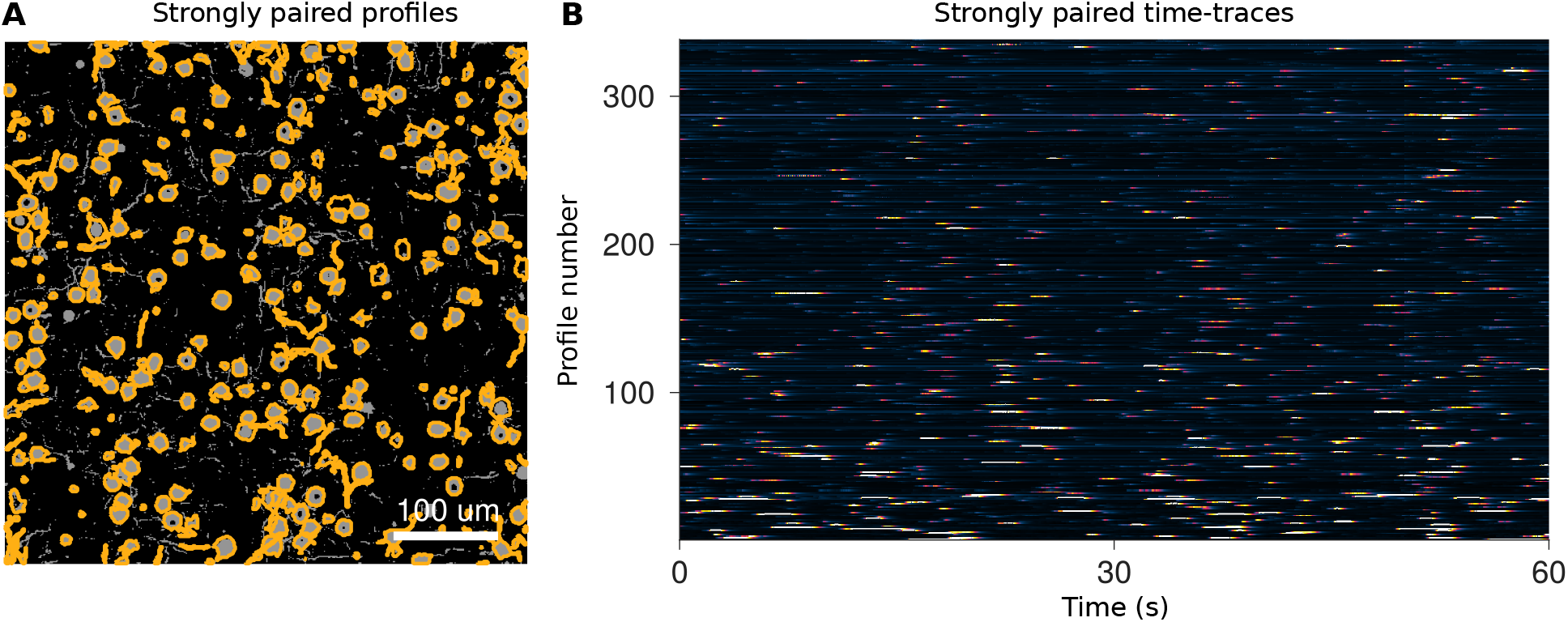
Results of using CNMF to analyze the simulated TPM video.

**Supplementary Figure 18:**
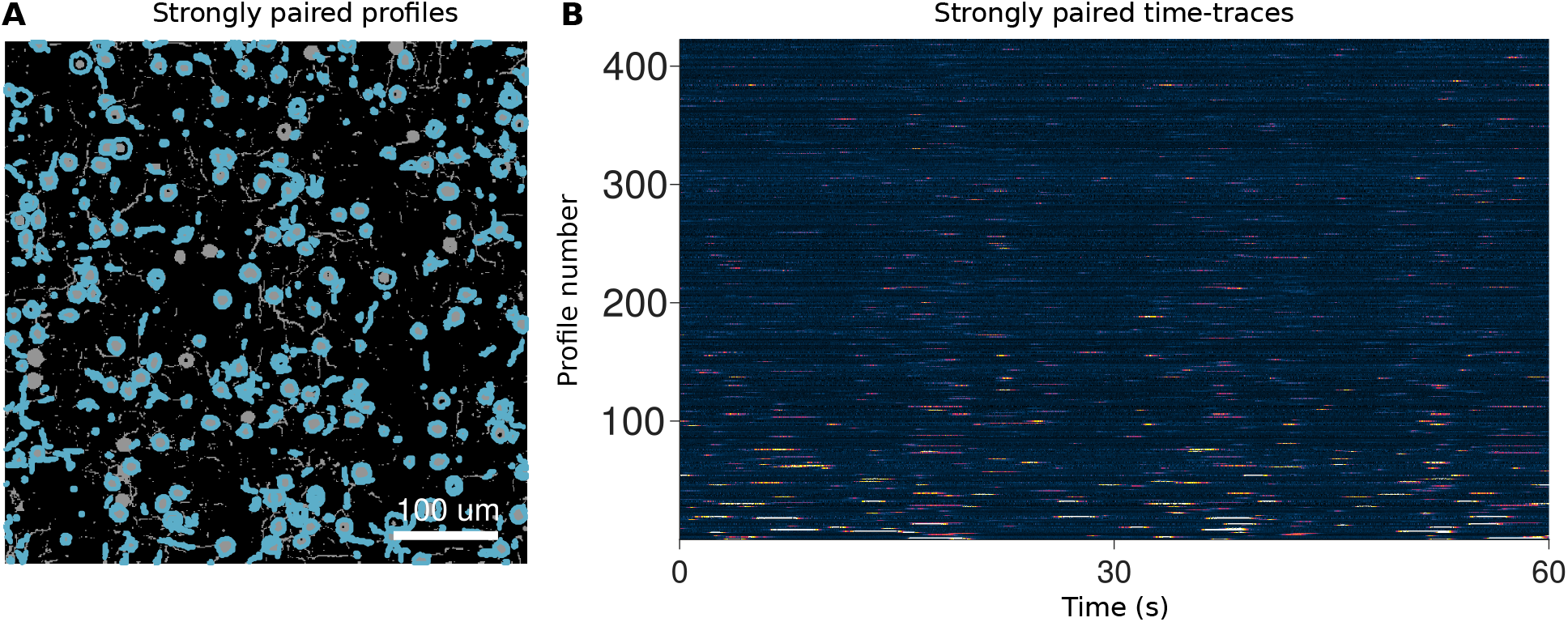
Results of using Suite2p to analyze the simulated TPM video.

**Supplementary Figure 19:**
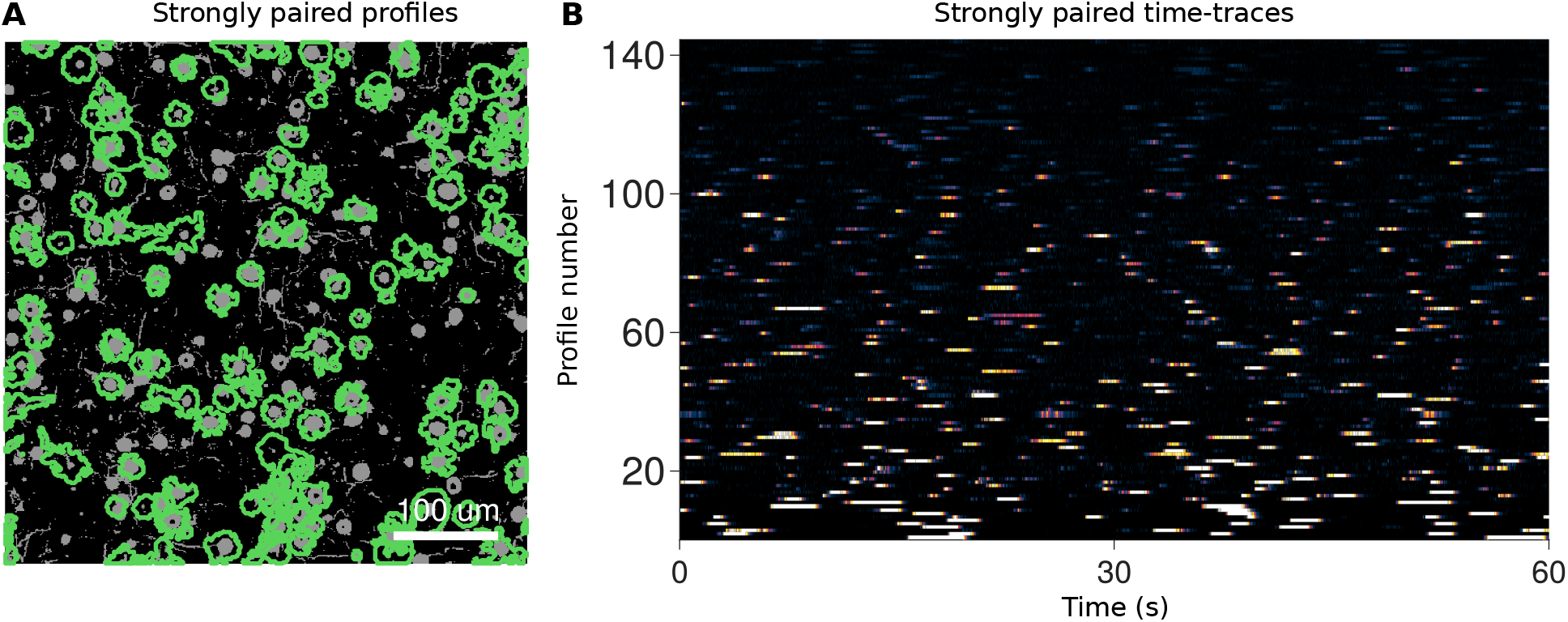
Results of using the PCA/ICA to analyze the simulated TPM video.

**Supplementary Figure 20:**
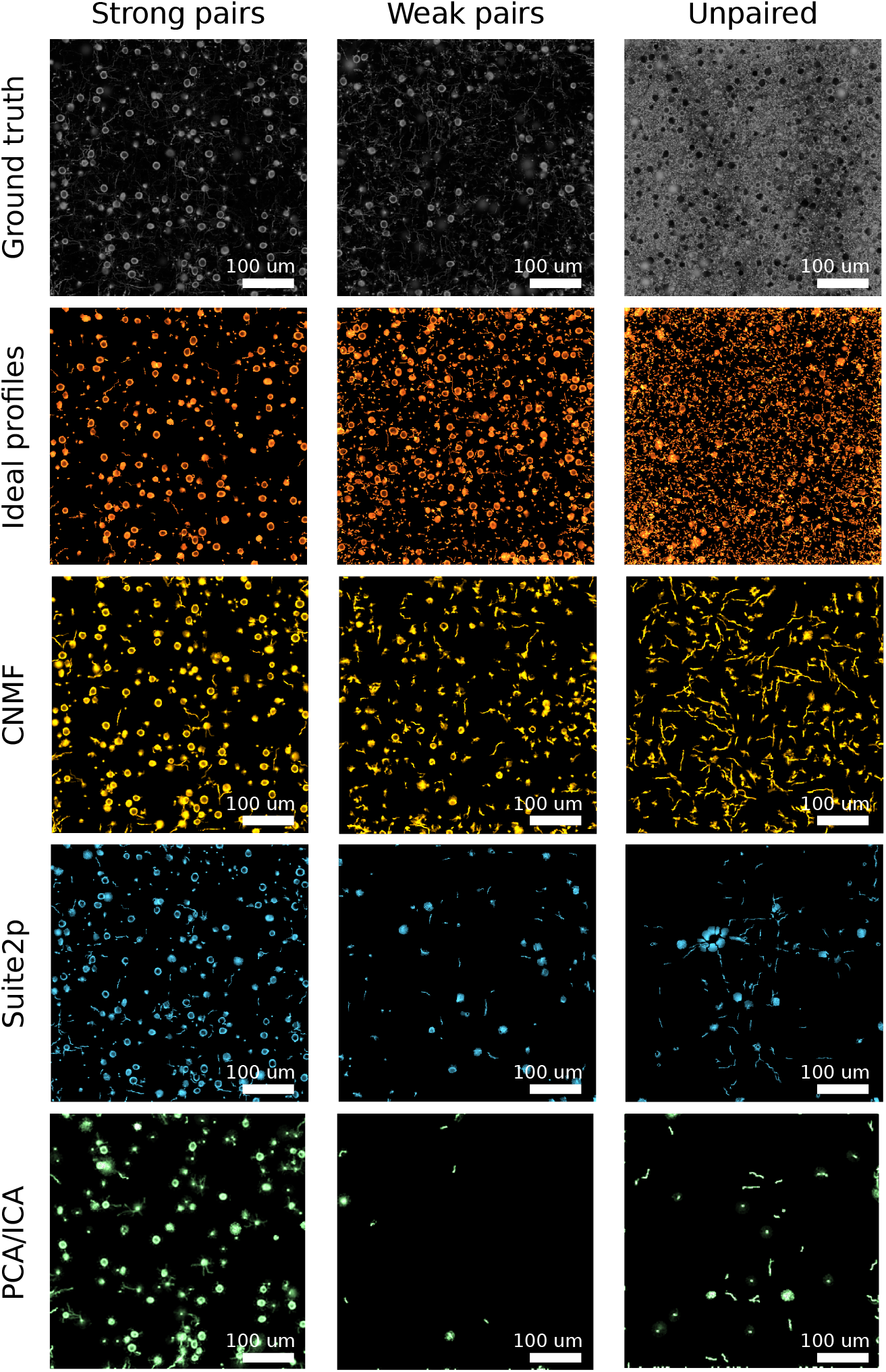
Profile shapes for strongly paired profiles (*ρ* > 0.5; left column), weakly paired profiles (0.1 < *ρ* < 0.5; middle column) and unpaired profiles (*ρ* < 0.1; right column). Most cells (top row), were unpaired. Paired cells tended to be found via their somatic signal. CNMF (second row) tended to find the most profiles and matched the most cells. CNMF, however, also found the most false-positives, which tended to be dendritic shapes. Suite2p (third row) found both fewer cells and fewer false positives. PCA/ICA had the lowest number of found cells but still had a significant number of false positives.

**Supplementary Figure 21:**
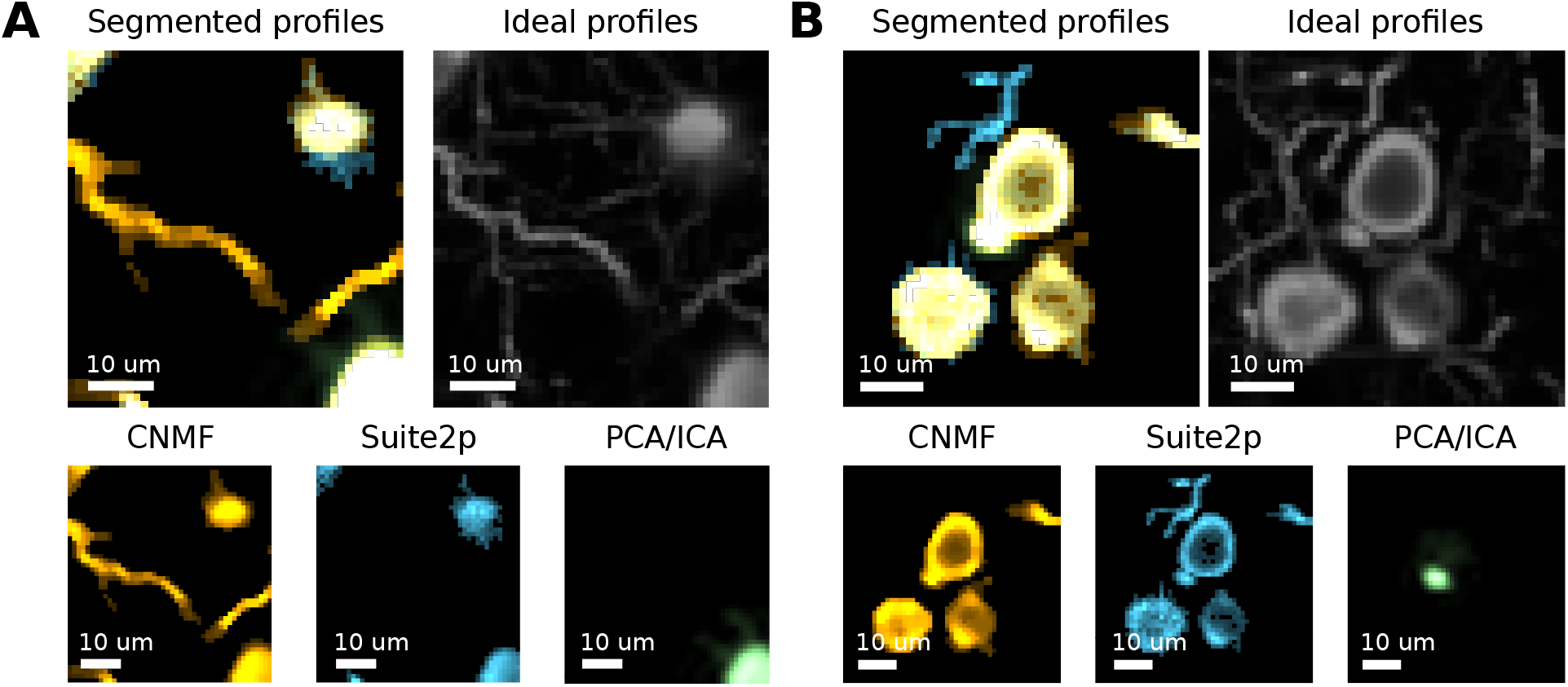
Examples of finer features in the segmented profiles. Amongst the profiles found by the three different methods (CNMF in red, Suite2p in blue and PCA/ICA in green) somas were often found by multiple methods while finer features, such as dendrites were often found by only one algorithm. A: A segment containing somas found by both CNMF and Suite2p (one also found using PCA/ICA) and a dendrite found only with CNMF. B: A segment containing somas found by both CNMF and Suite2p (one also found using PCA/ICA) and a dendrite found only with Suite2p.

**Supplementary Figure 22:**
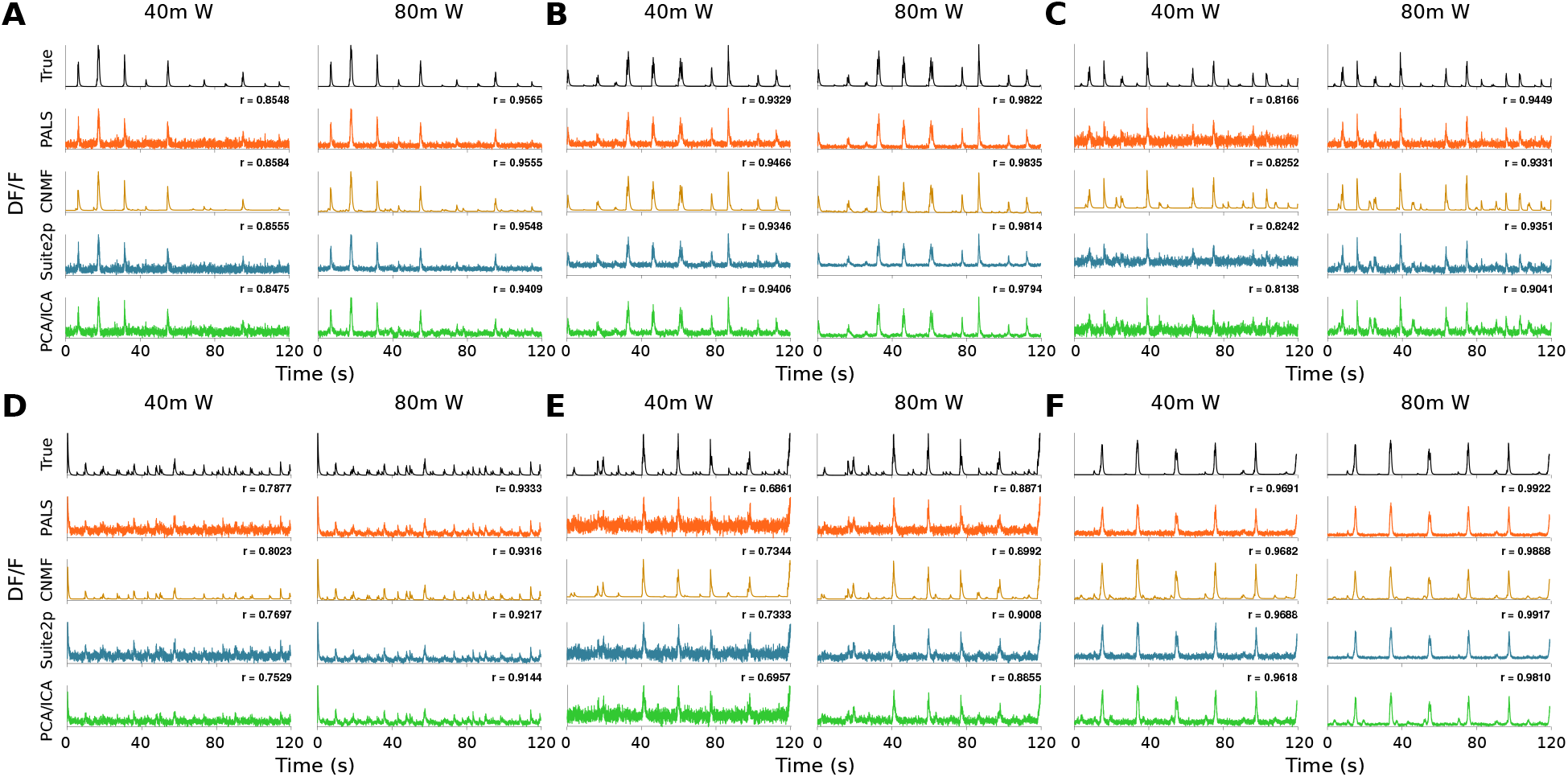
Examples of strongly paired time-traces in simulations using 0.6-NA Gaussian beams 40m W and 80m W power.

**Supplementary Figure 23:**
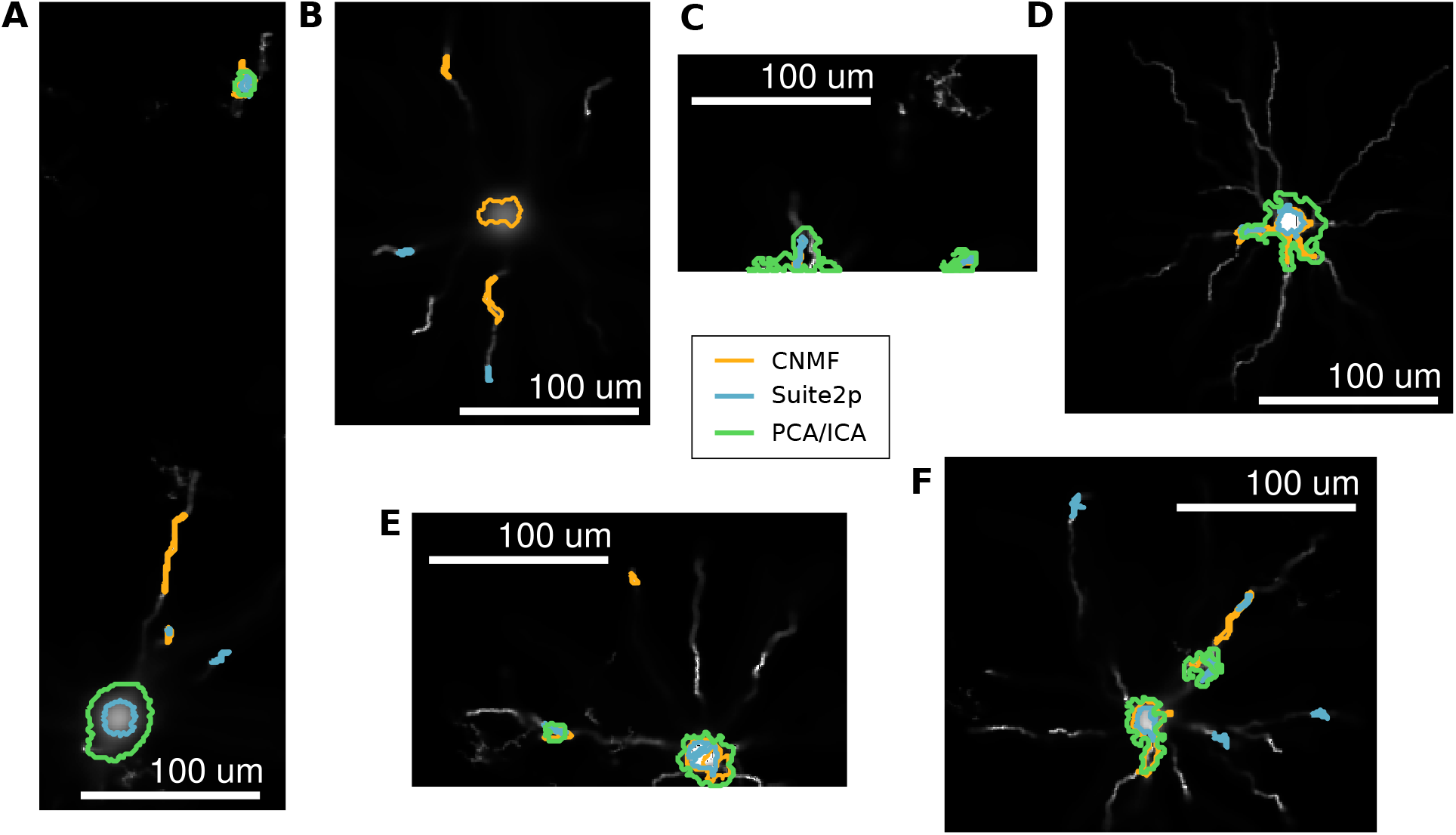
Examples of cells that were found with multiplicity by one or more algorithms. A: Such cells can have profiles that are hundreds of *μ*m away. B. More than two profiles can represent the same cell and often capture dendritic portions away from the soma. C: Such duplicity can happen when the F.O.V. cuts off a portion of the cell. D: Some examples were observed where the dendrite profile and soma profile were very close, where the profiles should have been merged. E: An example where two profiles represent the same soma and overlap. F: All three algorithms are susceptible to the multiplicity effect, proportionally to the number of profiles found.

**Supplementary Figure 24:**
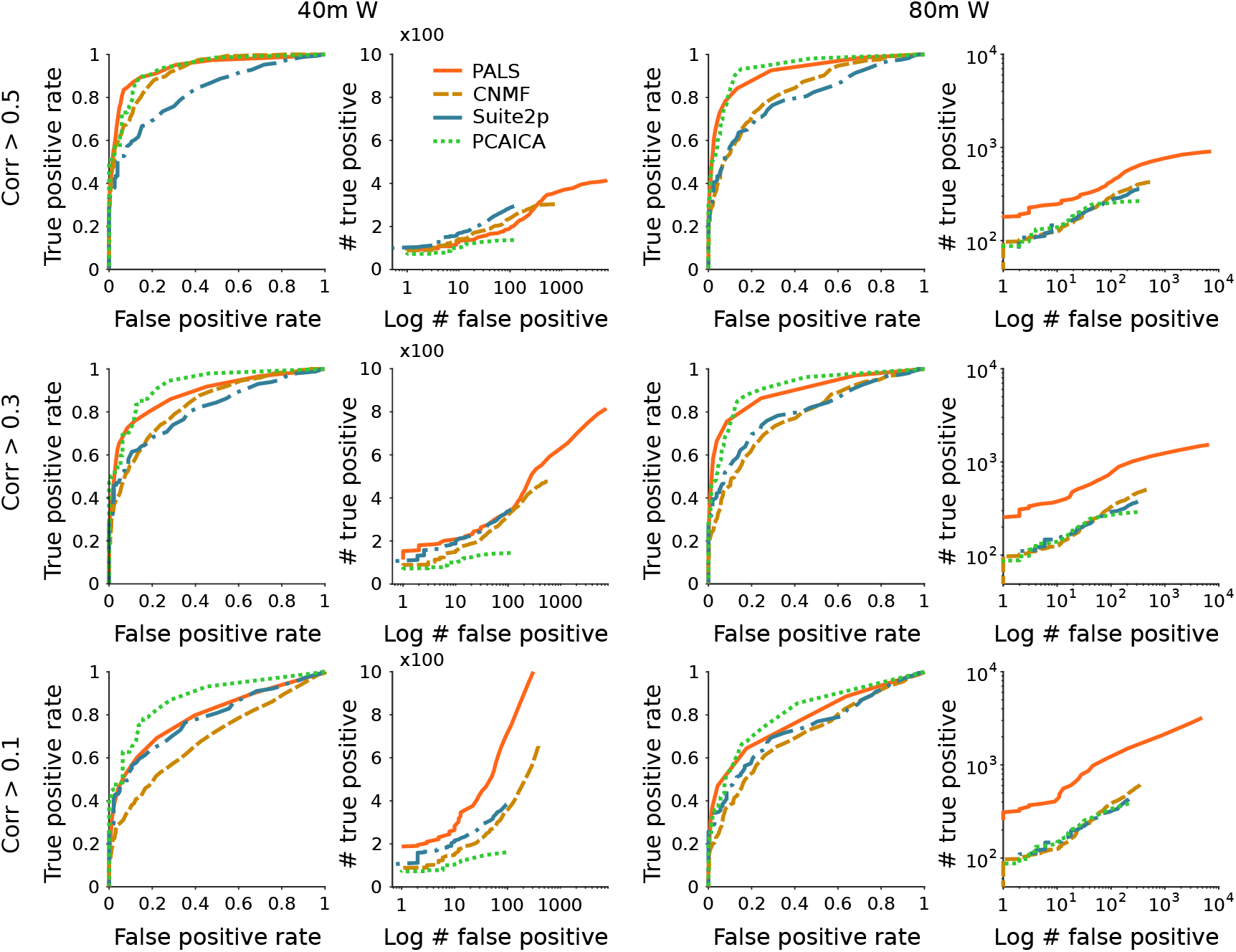
Examples ROC curves at different power levels (40 mW on the left and 80 mW on the right) and with different correlation cutoffs for determining true vs. artifact sources (0.1, 0.3, and 0.5 in the three rows). Each set of curves show on the left the rates of true vs. false, as a fraction of the total number of true and artifact cells found by each algorithm, and on the right the total number of true and artifact sources.

**Supplementary Figure 25:**
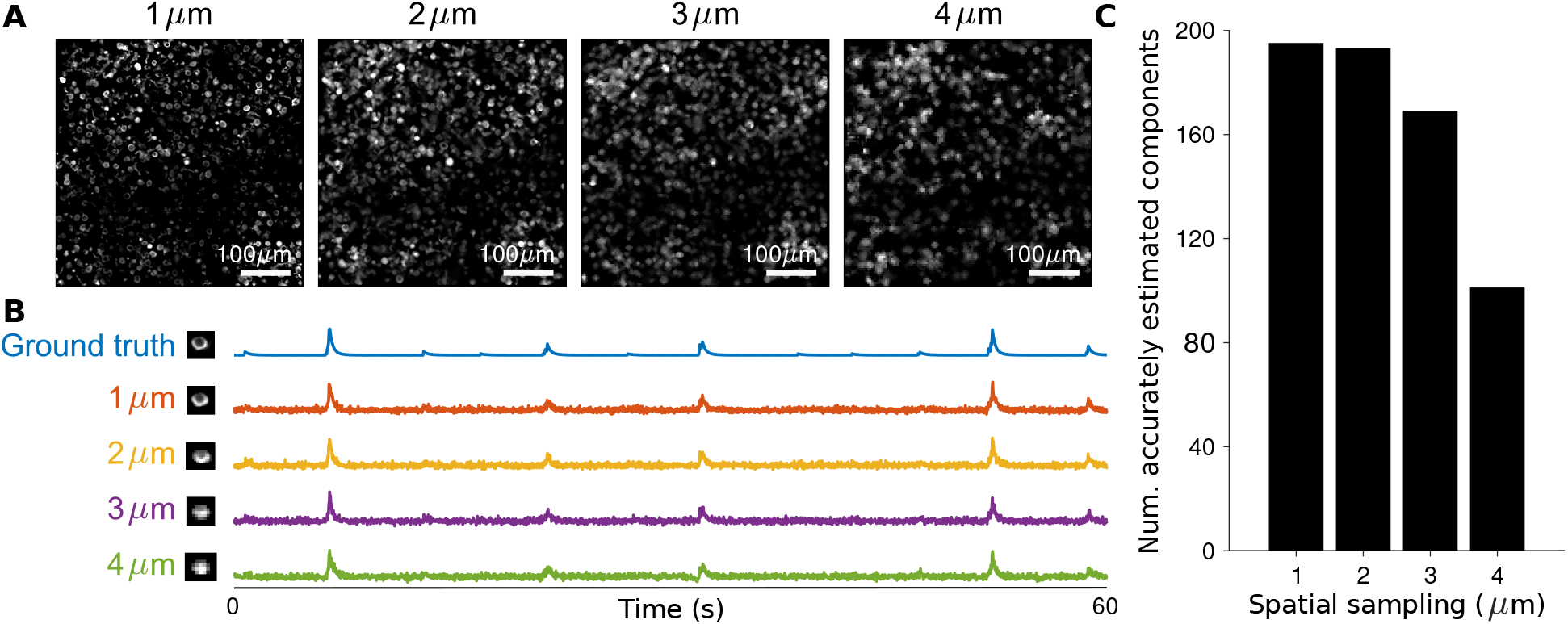
Analysis of signal loss as a function of subsampling interval. A: Mean images calculated from imaging movies of the same volume and activity, taken at 1, 2, 3, and 4*μ* m intervals. B: Example cell spatial profile and time traces at all four image subsampling resolutions, as compared to the ground truth used to generate the data. C: Comparison of the number of cells confidently found (correlation > 0.5) at all four resolutions indicate a steep fall-off after 3*μ* m subsampling.

**Supplementary Figure 26:**
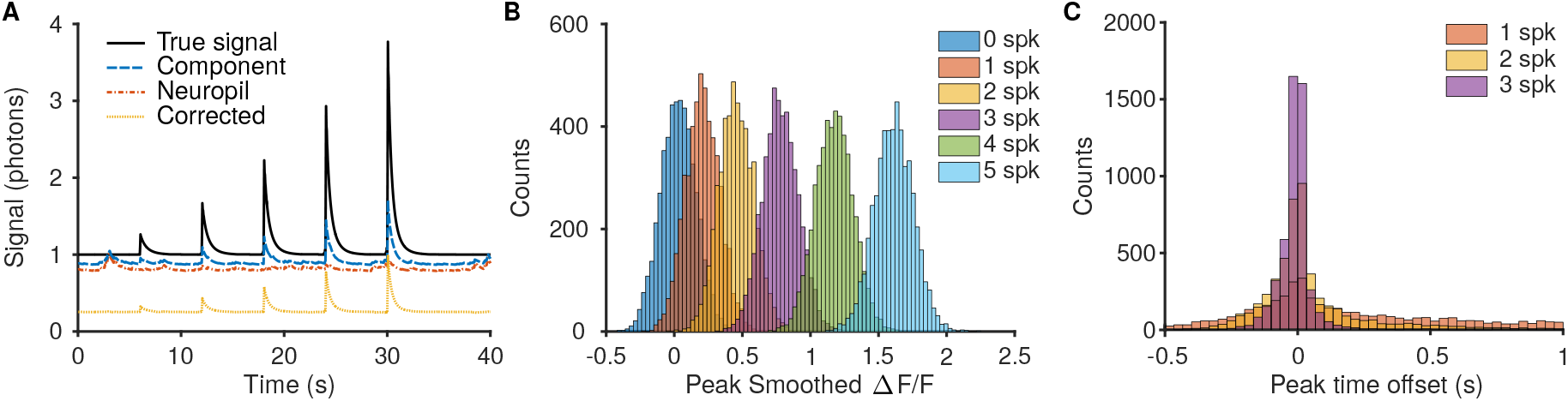
Comparison of resulting ΔF/F distinguishability for bursts of different spike numbers. A: Simulations were run by generating videos with preset spiking activity that ramped up the number of spikes in each burst. Much of the error in recovered component time-traces can be attributed to insufficiently removed neuropil, and using algorithmic techniques such as in Suite2P [18] can increase the accuracy of the time traces. B: The recovered ΔF/F for larger bursts was more reliable and more separable between the different burst strengths. For example there is less overlap between the peak ΔF/F for 5- and 4-spike bursts then for 1- and 2-spike bursts. C: The peak ΔF/F offset time is also more reliable for bursts with more spikes.

**Supplementary Figure 27:**
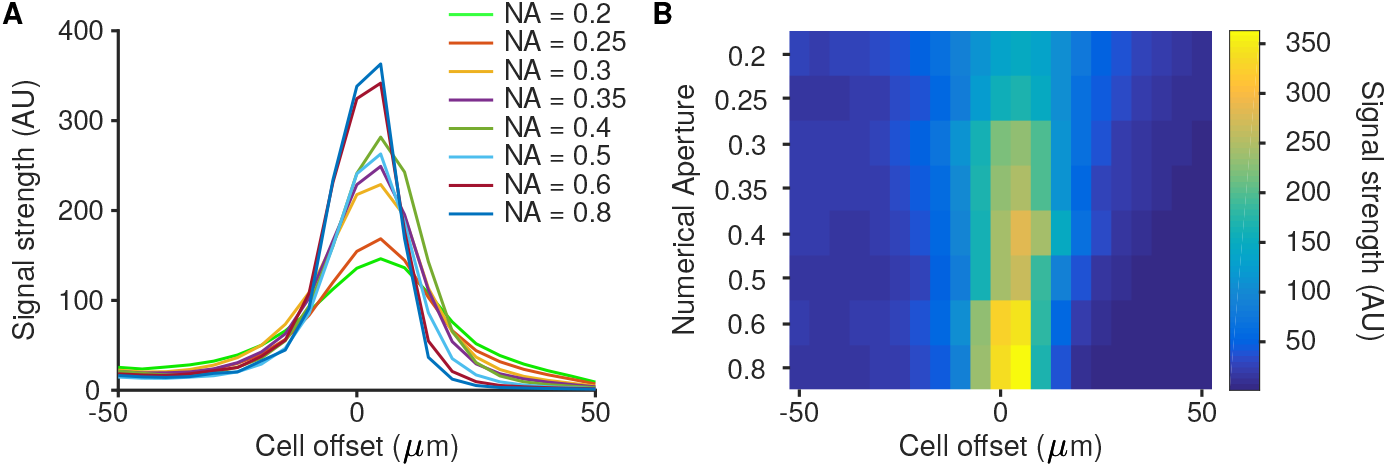
Analysis of signal strength as a function of numerical aperture (NA). A: Signal strength drops off as a function of the displacement of the cell from the peak PSF excitation. B: Same data plotted as an image to better compare spread and peak intensity of the various curves.

## Footnotes

1 Code will be made available post-review. Please contact A. Charles and A. Song for beta testing requests.

